# Comparing spatial null models for brain maps

**DOI:** 10.1101/2020.08.13.249797

**Authors:** Ross D. Markello, Bratislav Misic

## Abstract

Technological and data sharing advances have led to a proliferation of high-resolution structural and functional maps of the brain. Modern neuroimaging research increasingly depends on identifying correspondences between the topographies of these maps; however, most standard methods for statistical inference fail to account for their spatial properties. Recently, multiple methods have been developed to generate null distributions that preserve the spatial autocorrelation of brain maps and yield more accurate statistical estimates. Here, we comprehensively assess the performance of ten published null frameworks in statistical analyses of neuroimaging data. To test the efficacy of these frameworks in situations with a known ground truth, we first apply them to a series of controlled simulations and examine the impact of data resolution and spatial autocorrelation on their family-wise error rates. Next, we use each framework with two empirical neuroimaging datasets, investigating their performance when testing (1) the correspondence between brain maps (e.g., correlating two activation maps) and (2) the spatial distribution of a feature within a partition (e.g., quantifying the specificity of an activation map within an intrinsic functional network). Finally, we investigate how differences in the implementation of these null models may impact their performance. In agreement with previous reports, we find that naive null models that do not preserve spatial autocorrelation consistently yield elevated false positive rates and unrealistically liberal statistical estimates. While spatially-constrained null models yielded more realistic, conservative estimates, even these frameworks suffer from inflated false positive rates and variable performance across analyses. Throughout our results, we observe minimal impact of parcellation and resolution on null model performance. Altogether, our findings highlight the need for continued development of statistically-rigorous methods for comparing brain maps. The present report provides a harmonised framework for benchmarking and comparing future advancements.

## INTRODUCTION

The brain is organized as a series of nested and increasingly multi-functional neural circuits. The connections and interactions among these circuits ultimately manifest as unique topographic distributions of structural and functional properties. Recent advances in imaging, tracing, and recording technologies (Insel et al. 2013), together with global data sharing initiatives (Casey et al. 2018, Poldrack et al. 2013, Sudlow et al. 2015, Van Essen et al. 2013), have resulted in the generation of high-resolution maps of many of these properties, including gene expression (Akbarian et al. 2015, Hawrylycz et al. 2012), cytology (Scholtens et al. 2018, von Economo and Koskinas 1925), receptor densities (Beliveau et al. 2017, Norgaard et al. 2020, Zilles and Amunts 2009, Zilles et al. 2004), intracortical myelin (Burt et al. 2018, Whitaker et al. 2016), and functional organization (Bellec et al. 2010, Damoiseaux et al. 2006, Margulies et al. 2016, Murray et al. 2014, Shafiei et al. 2020, Yeo et al. 2011).

Increasingly, modern scientific discovery in neuroimaging research involves identifying correspondences between the topographies of brain maps (Baum et al. 2020, Demirtaş et al. 2019, Gao et al. 2020, Hansen et al. 2020, Shafiei et al. 2020, Vázquez-Rodríguez et al. 2019, Wang et al. 2019); however, standard methods for statistical inference fall short when making such comparisons (Alexander-Bloch et al. 2013, 2018, Breakspear et al. 2004, Burt et al. 2020, Fulcher et al. 2020, Gordon et al. 2016). Namely, in spatially-embedded systems—like the brain—neighboring data points are not statistically independent, violating the assumptions of many common inferential frameworks. As an example, consider computing a correlation between two brain maps. When using a standard parametric null (i.e., the Student’s *t*-distribution), the spatial autocorrelation of the maps violates the inherent requirement that the model errors are independent and identically distributed (*i.i.d.*). When using a standard non-parametric null (i.e., random permutations of one of the feature maps), the spatial auto-correlation violates the requirement of exchangeability. In both instances, the calculated *p*-value will be inflated, yielding increased family-wise error rates (FWER, or type 1 error rates) across analyses.

The impact of spatial autocorrelation on statistical inference has long been known in fields like geostatistics and ecology (Cliff and Ord 1970, Legendre 1993), and there have been significant developments in these fields to account for and overcome its influence (Cressie 2015, Deblauwe et al. 2012, Dray 2011, Dutilleul et al. 1993, Fortin and Jacquez 2000, Wagner and Dray 2015). Acknowledgement of the impact of spatial autocorrelation in neuroimaging is far more recent. Only in the past several years have frameworks been popularized to overcome the shortcomings of standard null models. These frameworks generate null distributions that preserve the spatial autocorrelation of brain maps, yielding statistical inferences that more accurately reflect the underlying data.

Spatially-constrained null models in neuroimaging fall into two broad families: non-parametric spatial permutation models (Alexander-Bloch et al. 2013, 2018, Gordon et al. 2016) and parameterized data models (Burt et al. 2018, 2020, Vos de Wael et al. 2020). In nonparametric spatial permutation models, the cortical surface is represented as a sphere to which random rotations are applied, generating surface maps with randomized topography but identical spatial autocorrelation. In parameterized data models, spatial autocorrelation is estimated from the empirical brain maps and used to generate surrogate nulls with randomized topography and similar—though not identical—spatial autocorrelation. Since their development, these models have been adapted by several researchers (Baum et al. 2020, Cornblath et al. 2020, Váša et al. 2018, Vázquez-Rodríguez et al. 2019). To our knowledge, there have been at least ten distinct implementations of null frameworks applied to statistical estimates of brain maps.

One of the earliest implementations of these null models, proposed in Alexander-Bloch et al. (2013) and later formalized in Alexander-Bloch et al. (2018), described a non-parametric method by which the cortical surface was subjected to spatial rotations. The principal challenge of implementing this method is that the medial wall—for which most brain maps contain no data—can be rotated into the cortical surface. This is an important consideration because the loss of data caused by this medial wall rotation can bias results. To address this problem, researchers have opted to either discard the missing data (Baum et al. 2020, Cornblath et al. 2020), assign the nearest data to missing parcels (Vázquez-Rodríguez et al. 2019), or ignore the medial wall entirely (Váša et al. 2018). Other groups have devised alternative methods that do not rely on spatial rotations but use generative models instead. These parameterized frameworks vary in their conceptualization and implementation of the data-generating process, ranging from a spatial lag model (Burt et al. 2018) to spectral randomization (Vos de Wael et al. 2020, Wagner and Dray 2015) to variogram matching (Burt et al. 2020). How these different models perform when applied to the same experimental questions and datasets remains unclear.

Here, we comprehensively compare how ten published null frameworks control the FWER in statistical analyses of neuroimaging data. First, using comprehensive simulations we examine the accuracy of statistical inferences drawn by each of the null models and their respective false positive rates. Next, relying on open-access empirical datasets (Neurosynth [Yarkoni et al. 2011]; Human Connectome Project [Van Essen et al. 2013]), we apply each null framework to two prototypical analyses: (1) assessing the correspondence between brain maps (e.g., correlating two activation maps) and (2) assessing the spatial distribution of a feature within a partition (e.g., quantifying the enrichment of T1w/T2w ratios in resting state networks). In all analyses we systematically examine the impact of parcellation—which differentially modifies the spatial structure of the underlying data—on the performance of the null frameworks.

## METHODS

### Code and data availability

All code used for data processing, analysis, and figure generation is available on GitHub (https://github.com/netneurolab/markello_spatialnulls) and directly relies on the following open-source Python packages: BrainSMASH (Burt et al. 2020), BrainSpace (Vos de Wael et al. 2020), IPython (Pérez and Granger 2007), Jupyter (Kluyver et al. 2016), Matplotlib (Hunter 2007), NeuroSynth (Yarkoni et al. 2011), NiBabel (Brett et al. 2020), NumPy (Oliphant 2006, Van Der Walt et al. 2011), Pandas (McKinney 2010), PySurfer (Waskom et al. 2020b), Scikit-learn (Pedregosa et al. 2011), SciPy (Virtanen et al. 2020), and Seaborn (Waskom et al. 2020a). Additional software used in the reported analyses includes FreeSurfer (v6.0.0, http://surfer.nmr.mgh.harvard.edu/; Fischl et al. 1999) and the Connectome Workbench (v1.4.2, https://www.humanconnectome.org/software/connectome-workbench; Marcus et al. 2011).

### Data

#### NeuroSynth association maps

To replicate the analyses described in Alexander-Bloch et al. (2018) we downloaded “association” maps from NeuroSynth (Yarkoni et al. 2011), a meta-analytic tool that synthesizes results from published MRI studies by linking voxel activation coordinates reported in an article to keywords in its abstract. We restricted the maps from NeuroSynth to terms that overlapped with those defined in the Cognitive Atlas, an ontological database for defining and relating cognitive processes (https://www.cognitiveatlas.org/concepts; Poldrack 2011, Poldrack et al. 2011, Poldrack and Yarkoni 2016; refer to Table S1 for a full list of terms). This resulted in 123 volumetric statistical maps which were then inflated to a mid-gray projection of FreeSurfer’s fsaverage5 surface using nearest neighbor interpolation.

#### Human Connectome Project

Group-averaged T1w/T2w (a proxy for intracortical myelin) data were downloaded from the S1200 release of the Human Connectome Project (HCP; Van Essen et al. 2013). To ensure consistency with the reported NeuroSynth analyses, data were resampled to FreeSurfer’s fsaverage5 surface following instructions on the Human Connectome Project wiki page (https://wiki.humanconnectome.org/display/PublicData/HCP+Users+FAQ).

#### Brain parcellations

In order to examine the impact of parcellations on the tested null models we used two multi-scale resolution atlases The first, referred to throughout the text as the “Cammoun atlas”, has five resolutions ranging from 68 to 1,000 parcels (68, 114, 219, 448, and 1,000 parcels), and was generated by dividing the Desikan-Killiany atlas (Desikan et al. 2006) into equally-sized sub-units based on group-level diffusion weighted imaging data (Cammoun et al. 2012). The second, referred to throughout the text as the “Schaefer atlas”, has ten resolutions ranging from 100 to 1,000 parcels in steps of 100 parcels (i.e., 100, 200, 300, and so on), and was generated via a gradient-weighted Markov Random Field model from resting state fMRI data (Schaefer et al. 2018).

NeuroSynth activation maps and the HCP T1w/T2w brain map were parcellated using all resolutions of the Schaefer and Cammoun atlases. Values for vertices lying on the medial wall of the surface mesh were not considered.

#### Network partitions

To investigate the partition specificity of brain maps we applied two common network definitions: the seven functional networks defined in Yeo et al. (2011) and the seven cytoarchitectonic classes proposed by von Economo and Koskinas (1925).

For the Cammoun atlas we used the fsaverage5 vertex-level Yeo-Krienen network assignments provided by FreeSurfer. To derive parcel-wise network assignments from this vertex representation we applied a winner-take-all approach; that is, the mode of the vertices in each parcel was used to select the representative network. For example, if a given parcel has twenty constituent vertices, ten of which are assigned to network A, three to network B, and seven to network C, a winner-take-all approach would assign that parcel to network A. For the Schaefer atlas we used the parcel assignments for the Yeo-Krienen networks provided by the original authors. To derive von Economo–Koskinas classes for parcels in both the Cammoun and Schaefer atlases we applied the classifiers provided by Scholtens et al. (2018) to the fsaverage5 surface, yielding vertex-level assignments. These assignments were then used with the winner-take-all approach to generate parcel-level network assignments.

### Null model frameworks

Here, we briefly describe null model frameworks as initially proposed in the literature, and then explain the specific details of the implementations used in the current report. For a summary overview of the implementations please refer to Table 1 and for an examination of their relative computational cost see Fig. S1.

**Table 1.** Null model frameworks. Overview of the null frameworks and the implementations used in the reported analyses. The *Description* column indicates the primary point of methodological divergence for each framework. The *Duplications* column indicates whether parcel values can be duplicated in the given framework. The *Citation* column indicates the first appearance of the described implementation in the neuroimaging literature, if known. Code and/or references to code for implementing all frameworks is available at https://github.com/netneurolab/markello_spatialnulls. Refer to Fig. S1 for comparisons of the computational time of each framework and Table S2 for an overview of the drawbacks of each null model.

| Naive models |  |  |  |
| --- | --- | --- | --- |
| *Method* | *Description* | *Duplications* | *Citation* |
| Parametric | Uses standard parametric distributions (e.g., Student’s *t*-distribution) | No | — |
| Non-parametric | Reorders parcels randomly, not taking into account their inherent spatial structure | No | — |
| Spatial permutation models |  |  |  |
| *Method* | *Description* | *Duplications* | *Citation* |
| Vázquez-Rodríguez | Assigns parcel centroid to closest rotated centroid based on minimum Euclidean distance | Yes | Vázquez-Rodríguez et al. (2019) |
| Baum | Projects parcel data to vertices, rotates, and takes the mode of vertices in each parcel | Yes | Baum et al. (2020) |
| Cornblath | Projects parcel data to vertices, rotates, and takes the mean of vertices in each parcel | Yes | Cornblath et al. (2020) |
| Váša | Uses an iterative procedure to uniquely assign parcels based on distance to the rotated parcels | No | Váša et al. (2018) |
| Hungarian | Uses the Hungarian algorithm to match original and rotated parcel centroids | No |  |
| Parameterized data models |  |  |  |
| *Method* | *Description* | *Duplications* | *Citation* |
| Burt-2018 | Uses a distance-dependent spatially autoregressive model to generate nulls | No | Burt et al. (2018) |
| Burt-2020 | Uses a variogram model to estimate spatial autocorrelation to generate nulls | No | Burt et al. (2020) |
| Moran | Uses a distance-dependent eigendecomposition to generate nulls | No | Vos de Wael et al. (2020) |

#### Naive models

Commonly used in the neuroimaging literature prior to the development of spatially-constrained frameworks, so-called naive models do not take into account the spatial structure of the data when constructing null distributions, commonly resulting in inflated family-wise error rates. We test two such naive models: a *parametric* and a *non-parametric* method.

##### Naive parametric method

Although the exact implementation of the parametric method varies based on the statistical test employed, all implementations share a reliance on standard null distributions. For example, when examining correlation values, the parametric method relies on the Student’s *t*-distribution; when examining z-statistics, this method uses the standard normal distribution.

##### Naive non-parametric method

The naive non-parametric approach uses a random permutation (i.e., reshuffling) of the data to construct a null distribution, destroying its inherent spatial structure. Each vertex or parcel is uniquely reassigned the value of another vertex or parcel for every permutation.

#### Spatial permutation models

First published in Alexander-Bloch et al. (2013) and later formalized in Alexander-Bloch et al. (2018), spatial permutation models generate spatially-constrained null distributions by applying random rotations to spherical projections of the brain. A rotation matrix (**R**) is applied to the three-dimensional coordinates of the brain (**V**) to generate a set of rotated coordinates (**V_rot_** = **VR**). The permutation is constructed by replacing the original values at each coordinate with those of the closest rotated coordinate. Rotations are generated independently for one hemisphere and then mirrored across the anterior-posterior axis for the other.

Application of the *Alexander-Bloch* rotation method to parcellated brain data entails several methodological choices, and in recent years at least four different adaptations on this technique have been published: Vázquez-Rodríguez et al. (2019), Váša et al. (2018), Baum et al. (2020), and Cornblath et al. (2020). We test all four of these adaptations, in addition to one other method (*Hungarian*) that we posit is a natural extension thereof. Note, however, that when analyzing vertex-level data all of these adaptations reduce to the original *Alexander-Bloch* technique.

##### Vázquez-Rodríguez method

The *Vázquez-Rodríguez* method, which serves as a direct adaptation of the original framework from Alexander-Bloch et al. (2018) but applied to parcellated brain data, was first used in Vázquez-Rodríguez et al. (2019). In this adaptation, vertex coordinates are replaced with those of parcel centroids. That is, a rotation is applied to the coordinates for the center-of-mass of each parcel, and parcels are reassigned the value of the closest rotated parcel (i.e., that with the minimum Euclidean distance). If the medial wall is rotated into a region of cortex the value of the nearest parcel is assigned instead, ensuring that all parcels have values for every rotation. This method for handling the medial wall consequently permits the duplicate reassignment of parcel values for every rotation, such that some parcel values may not be present in a given rotation and others may appear more than once. Note that the exact method used to define parcel centroids may impact the performance of this model (see *Methods: Null model implementation variability*).

##### Baum method

Used initially in Baum et al. (2020), this method projects parcellated brain data to a high-resolution surface mesh, assigning identical values to all the vertices within a given parcel. The projected mesh is subjected to the original spatial permutation reassignment procedure (i.e., from Alexander-Bloch et al. 2018) and re-parcellated by taking the modal (i.e., the most common) value of the vertices in each parcel. When the rotated medial wall completely subsumes a cortical parcel that region is assigned a value of NaN and is removed from subsequent analyses. Notably, this method can result in duplicate assignment of parcel values in each permutation.

##### Cornblath method

In this method implemented by Cornblath et al. (2020), parcellated data are projected to a high-resolution spherical surface mesh, rotated, and re-parcellated by taking the average (i.e., the arithmetic mean) of the vertices in each parcel. When the rotated medial wall completely subsumes a cortical parcel that region is assigned a value of NaN and is removed from subsequent analyses. Because the data are re-parcellated the likelihood of duplicated assignments is very low (though not exactly zero); however, the distribution of re-parcellated values will be slightly different than the original data distribution.

##### Váša method

The first known application of spatial permutations to parcellated data, the *Váša* method (Váša et al. 2018) attempted to resolve one of the primary drawbacks of the Alexander-Bloch method: duplicate reassignment of values. That is, this method was created so as to yield a “perfect” permutation of the original data for every rotation. Similar to the *Vázquez-Rodríguez* method, parcel centroids are used instead of vertex coordinates. In order to avoid duplicate reassignments, parcels are iteratively assigned by (1) finding the closest rotated parcel to each original parcel, and (2) assigning the most distant pair of parcels. This two-step process is then repeated for all remaining unassigned parcels until each has been reassigned. Parcels are reassigned without consideration for the medial wall or its rotated location. Note that the exact method used to define parcel centroids may impact the performance of this model (see *Methods: Null model implementation variability*).

##### Hungarian method

Similar to the *Váša* method, the *Hungarian* method attempts to uniquely reassign each parcel for every rotation. Instead of using an iterative process, however, which can result in globally sub-optimal assignments, this method uses the Hungarian algorithm to solve a linear sum assignment problem (Kuhn 1955). This method attempts to uniquely reassign each parcel such that the global reassignment cost is minimized, where cost is quantified as the distance between the original and rotated parcel centroid coordinates. The medial wall is ignored in all rotations and the optimal reassignment is determined without consideration for its location. Note that the exact method used to define parcel centroids may impact the performance of this model (see *Methods: Null model implementation variability*).

##### Null model implementations

The spherical projection of the fsaverage5 cortical mesh from FreeSurfer was used to define coordinates for all of the spatial permutation null models (Fischl et al. 1999). When parcel centroids were required for a model we used the procedure described in Vázquez-Rodríguez et al. (2019), *surface-projection averaging*, which includes: (1) calculating the arithmetic mean of the coordinates of all the vertices within a given parcel, and (2) using the coordinate of the surface vertex closest to this average (where closest is defined as minimizing Euclidean distance) to represent the parcel centroid. Other means of parcel centroid definition yielded similar results (see *Variability in parcel centroid definition*).

#### Parameterized data models

Distinct from the formulation of spatial permutation models proposed by Alexander-Bloch et al. (2018), parameterized data models do not rely on rotations to generate null distributions. Instead, these models generate surrogate null maps that retain spatial features characteristic of the data from which they are estimated. We test three such models, initially proposed for use in the neuroimaging (Burt et al. 2018, 2020) and ecology (Wagner and Dray 2015) literature.

##### Burt-2018 method

Described in Burt et al. (2018), this framework uses a spatial autoregressive model of the form **y** = *ρ* **Wy** to generate surrogate data. Here, **y** refers to a Box-Cox transformed, mean-centered brain feature of interest (i.e., a brain map), **W** is a weight matrix (derived from **D**, a matrix of the distance between brain regions, and *d*_0_, a spatial autocorrelation factor), and *ρ* is a spatial lag parameter. The parameters *ρ* and *d*_0_ are derived from the data via a least-squares optimization procedure and their estimates 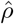 are 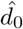 used to generate surrogate brain maps according to 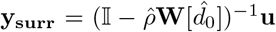, where 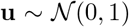 is a vector of random Gaussian noise. Rank-ordered values in the **y_surr_** map are replaced with corresponding values from the original **y**.

##### Burt-2020 method

Two years after introducing their spatial autoregressive method, Burt et al. (2020) proposed a novel model to generate surrogate data using variogram estimation. The method operates in two main steps: (1) randomly permute the values in a given brain map, and (2) smooth and re-scale the permuted values to reintroduce spatial autocorrelation characteristic of the original, non-permuted data. Reintroduction of spatial autocorrelation onto the permuted data is achieved via the transformation 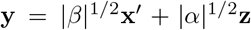, where **x′** is the permuted data, 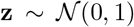 is a vector of random Gaussian noise, and *α* and *β* are estimated via a least-squares optimization between variograms of the original and permuted data. When applied to empirical data, rank-ordered values in the surrogate map are replaced with corresponding values from the original brain map; surrogates maps generated from simulated data use the raw values of **y**.

##### Moran method

Originally developed in the ecology literature (Dray 2011, Wagner and Dray 2015), Moran spectral randomization (MSR) has only been recently applied to neuroimaging data (Paquola et al. 2020, Royer et al. 2020, Vos de Wael et al. 2020). Similar to the other parameterized data methods, MSR principally relies on a spatially-informed weight matrix **W**, usually taking the form of an inverse distance matrix between brain regions. However, rather than using **W** to estimate parameters via a least-squares approach, MSR uses an eigende-composition of **W** to compute spatial eigenvectors that provide an estimate of autocorrelation. These eigenvectors are then used to impose a similar spatial structure on random, normally distributed surrogate data.

##### Null model implementations

All parameterized data models require an input distance or weight matrix providing information about the spatial structure of the corresponding brain maps. In the present report we calculated vertex-vertex surface distance matrices on the pial surface of the fsaverage5 cortical mesh from FreeSurfer. Shortest paths on the surface were not allowed to traverse the medial wall (see *Methods: Geodesic distances along the medial wall*). Parcel-parcel distance matrices were calculated for each parcellation and parcellation resolution by averaging the distance between every vertex in two parcels. The parcel-parcel (or vertex-vertex) distance matrix was used without modification for the *Burt-2018* and *Burt-2020* methods; the inverse of the distance matrix—with the diagonal set to 1—was used as an input for the *Moran* method. Although the use of the full, non-sparsified distance matrix for the *Moran* method differs somewhat from its traditional application in neuroimaging research (Paquola et al. 2020, Royer et al. 2020, Vos de Wael et al. 2020), we believe the methodology used in the current report should yield better-matched surrogates. Finally, note that the *Burt-2020* method is unique amongst all the spatial parameterized data models in that it has several hyper-parameters that can be modified to control its performance. We use the default parameters provided by the BrainSMASH software (Burt et al. 2020), with the exception of those analyses of vertex-level data where we modify the knn parameter (see Fig. S2).

### Simulated data analyses

To examine the efficacy of the null frameworks we performed a series of controlled simulations with a known “ground truth”. Directly adapting methodology used in Burt et al. (2020), we derived pairs of brain maps with a specified Pearson correlation (target *r* = 0.015 ± 0.005) from spatially-autocorrelated Gaussian random fields (GRFs). To generate each pair of brain maps we started by creating a pair of dense, three-dimensional GRFs. The spatial autocorrelation of the GRFs was varied across seven different levels by modifying the slope of the field’s power spectral density (*α* = 0–3.0 in increments of 0.5, where zero indicates random Gaussian noise; see Appendix 2 of Burt et al. (2020) for information on the mathematical foundation of this approach). The pair of GRFs were projected to the fsaverage5 cortical mesh using FreeSurfer’s `mri_vol2surf`, the medial wall was removed (i.e., set to NaN), and the resulting brain maps were correlated to ensure that they fell within the target correlation limits. We generated 1,000 pairs of brain maps at each level of spatial autocorrelation, resulting in 7,000 total pairs of maps with correlations in the range 0.145–0.155. These data were also parcellated with the Cammoun and Schaefer atlases to yield lower-resolution representations.

To examine the comparative efficacy of the null models we applied each model to the simulated brain maps. For one map in each pair we used the given null method to generate 1,000 null maps, which we then correlated with the other map to yield a null distribution of correlation coefficients. We used this null distribution to estimate the two-tailed *p*-value for the original correlation between the pair of maps. This process was repeated across all levels of spatial autocorrelation (*α*) for each null framework, and was performed for the simulations at both the vertex-level fsaverage5 resolution and all Cammoun and Schaefer parcellation resolutions. We examined the generated *p*-values as a function of the spatial autocorrelation, *α*, of the simulated maps across all null frameworks.

Next, we sought to assess the false positive rate (FPR) of each of the null frameworks. If we examine a set of randomly-correlated pairs of brain maps, each null framework should yield a FPR approximately equal to the chosen statistical alpha (i.e., a 5% FPR when using an alpha of *p* ≤ 0.05). To generate these sets of randomly-correlated brain map pairs we shuffled the brain map pairs from the previous analysis, destroying the original correlation structure. As in the previous analysis, for one map in each pair we used each null framework to generate 1,000 null maps, which we then correlated with the other map in the pair to yield a null distribution of correlation coefficients. We estimated the two-tailed *p*-value of the original correlation, repeated this process for all brain map pairs across all levels of spatial autocorrelation (*α*) for each null framework, and from all these *p*-values we estimated the probability at which a given null framework would yield a significant test at the *p* ≤ 0.05 threshold. We examined the generated probabilities as a function of spatial autocorrelation across all null frameworks.

Note that for vertex-wise representations of the data all of the spatial permutation null models reduce to the original method proposed by Alexander-Bloch et al. (2018); however, where applicable we retain the *Vázquez-Rodríguez* label for consistency with results from the parcellated data. Finally, to ensure comparability across spatial permutation models, the same angular rotations were used across frameworks (e.g., for *Vázquez-Rodríguez*, *Váša*, and *Hungarian*).

### Empirical data analyses

Following the analyses of simulated data, we examined the performance of the null frameworks on two empirical neuroimaging datasets. Here, our choice of datasets and associated analytic procedures were designed to replicate and extend analyses originally reported in Alexander-Bloch et al. (2018) and Burt et al. (2020). We use (1) data from NeuroSynth to examine the correspondence between multiple brain maps (as in Alexander-Bloch et al. 2018), and (2) data from HCP to examine whether the spatial distribution of a brain map is circumscribed by predefined partitions (e.g., intrinsic networks, cytoarchitectonic classes; as in Burt et al. 2020). Null distributions for each framework were constructed from 10,000 permutations, rotations, or surrogates, and when applicable, the same angular rotations were used across spatial permutation models (e.g., for *Vázquez-Rodríguez*, *Váša*, and *Hungarian*). We analyze only parcellated data here to allow for better disambiguation between all ten null models.

#### Testing correspondence between brain maps (NeuroSynth)

Following procedures outlined by Alexander-Bloch et al. (2018), NeuroSynth maps were correlated to generate a term × term (123 × 123) correlation matrix, indicating the extent to which pairs of cognitive terms share a similar spatial pattern; instead of using vertex-wise values as in the original publication, correlation matrices were generated from parcellated data (see *Methods: Brain parcellations*). We applied each of the ten null frameworks to examine which of the resulting 7,503 unique correlations were significant. For each framework the NeuroSynth maps were permuted (or rotated, or used to generate surrogate data, as appropriate) and re-correlated, yielding a 123 × 123 null correlation matrix for each permutation representing the correlations between the original and null term maps. The largest absolute value (excluding the diagonal) of each permuted correlation matrix was retained and stored, generating a null distribution of 10,000 correlations; this procedure provides family-wise control for multiple comparisons (Alexander-Bloch et al. 2018, Westfall and Young 1993). We used this null distribution to estimate the two-tailed *p*-values for the correlations in the original term-by-term matrix, thresholding the matrix at *p* ≤ 0.05.

For the naive parametric null method, we used the Student’s *t*-distribution to generate *p*-values for each correlation between cognitive maps. For the parameterized null frameworks where surrogate data depend on the input brain maps, we constructed 10,000 surrogate maps separately for each cognitive term; to maintain consistency we used the same 10,000 random data vectors (e.g., **u** in Burt-2018, **z** in Burt-2020) for every term.

#### Testing partition specificity (HCP)

Using parcellated T1w/T2w data (Van Essen et al. 2013), we calculated the average value of all parcels within each of seven intrinsic functional networks (Yeo et al. 2011) and seven cytoarchitectonic classes (Scholtens et al. 2018, von Economo and Koskinas 1925). We then applied each of the ten null frameworks to examine which of these averages were significantly higher or lower than would be otherwise expected. For each framework, the parcel values were permuted (or rotated, or used to generate surrogate data, as appropriate) and re-averaged within the partitions, yielding a distribution of 10,000 null values for each network or class. We used these null distributions to estimate the two-tailed *p*-values for the original partition averages at *α* = 0.05.

For the naive parametric null framework we used the Student’s *t*-distribution to generate *p*-values, testing the distribution of parcel values for each network against zero, where the overall distribution of parcel values were z-scored prior to segregation into networks.

### Null model implementation variability

While the present report is primarily interested in how the different null models compare to one another, we also wanted to investigate the extent to which methodological choices within each model impact their performance. That is, most of the null models require researchers to make certain decisions when implementing them in practice. We investigate two such choices: (1) definition of parcel centroids for the spatial permutation nulls models, and (2) definition of the geodesic distance matrix for parameterized data models.

#### Variability in parcel centroid definition

We examined the impact of parcel centroid definition on three spatial permutation null models: *Vázquez-Rodríguez*, *Váša*, and *Hungarian*. Parcel centroids were defined using three different techniques operating on the spherical representation of the fsaverage5 surface (Fischl et al. 1999): (1) averaging, (2) averaging with surface-projection, and (3) geodesic distance minimization. In *averaging*, the coordinates for all vertices belonging to a parcel were averaged and used to represent the parcel centroid without further modification (Váša et al. 2018). Because the vertices are defined o n a sphere, these averaged-coordinate centroids will always fall beneath the surface of the cortical mesh. To resolve this, *averaging with surface-projection* performs the same procedure as *averaging* but then selects the coordinates of the closest vertex on the surface of the cortical mesh to represent the parcel centroid (Vázquez-Rodríguez et al. 2019). In the case of oblong or C-shaped parcels, however, surface-projected centroids may still fall outside the parcel boundaries. An alternative approach, *geodesic distance minimization*, avoids this shortcoming by computing a vertex-by-vertex geodesic distance matrix separately for each parcel. Each distance matrix is averaged across columns and the coordinates of the vertex with the smallest average is used to represent the parcel centroid. Parcellating data with non-spatially contiguous atlases (e.g., the Yeo-Krienen functional networks; Yeo et al. 2011) is not recommended when defining parcel centroids for use with spatial permutation null models.

We generated ten sample reassignments for all nine combinations of parcel centroid definition method and the three aforementioned spatial permutation null models. The normalized Hamming distance was used to compare the similarity between all generated reassignments (Hamming 1950), which was computed as the proportion of those vector elements between any two reassignments which disagree:

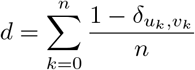

where *u* and *v* are any two reassignments of length *n* and 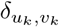 is the Kronecker delta function which is 1 when *u_k_* = *v_k_* and 0 otherwise.

#### Geodesic distances along the medial wall

We also investigated the degree to which constraining calculation of surface-based distance matrices to disallow travel along the medial wall impacts the outcomes of parameterized data models. We generated two distance matrices for each parcellation and resolution, either (1) permitting or (2) disallowing the travelled paths to cross the medial wall. Surface distance between vertices was calculated using Dijkstra’s algorithm on a graph representing the pial cortical mesh of the fsaverage5 surface (Fischl et al. 1999). Parcel-to-parcel distance was calculated as the average distance between every surface vertex in two parcels.

We used the generated distance matrices to create 1,000 surrogates for each parameterized data method, yielding 6,000 total surrogates (two distance matrices times three parameterized methods), and assessed the similarity of the resulting surrogates within each method using linear correlations. Statistics for the correlations were calculated by first transforming the correlations using the Fisher z-transform, estimating (1) the mean and (2) the 2.5 and 97.5% CI of the transformed estimates (via 10,000 bootstraps), and then applying the inverse Fisher z-transform to these estimates. As the parameterized data models require an input brain map we used the parcellated T1w/T2w data to generate these surrogates.

## RESULTS

We performed four analyses to investigate the impact of ten null models on controlling the FWER in statistical assessments of brain data.

### Null model performance on simulated brain maps

To quantitatively assess how the null models perform on data with a known “ground truth” we performed a series of controlled simulations. First, directly adapting methodology from Burt et al. (2020), we simulated 1,000 pairs of correlated brain maps (*r* = 0.15) derived from Gaussian random fields (GRFs) across seven different degrees of spatial autocorrelation (*α*; Fig. 2a,b). We applied each null framework to every pair of brain maps to generate 1,000 null maps per pair, yielding one million null maps per null framework per level of spatial autocorrelation (although we find that the exact number of null maps does not meaningfully influence our results, Fig. S3). We used these nulls to generate a two-tailed *p*-value on the original correlation between each brain map pair.

**Figure 1.**
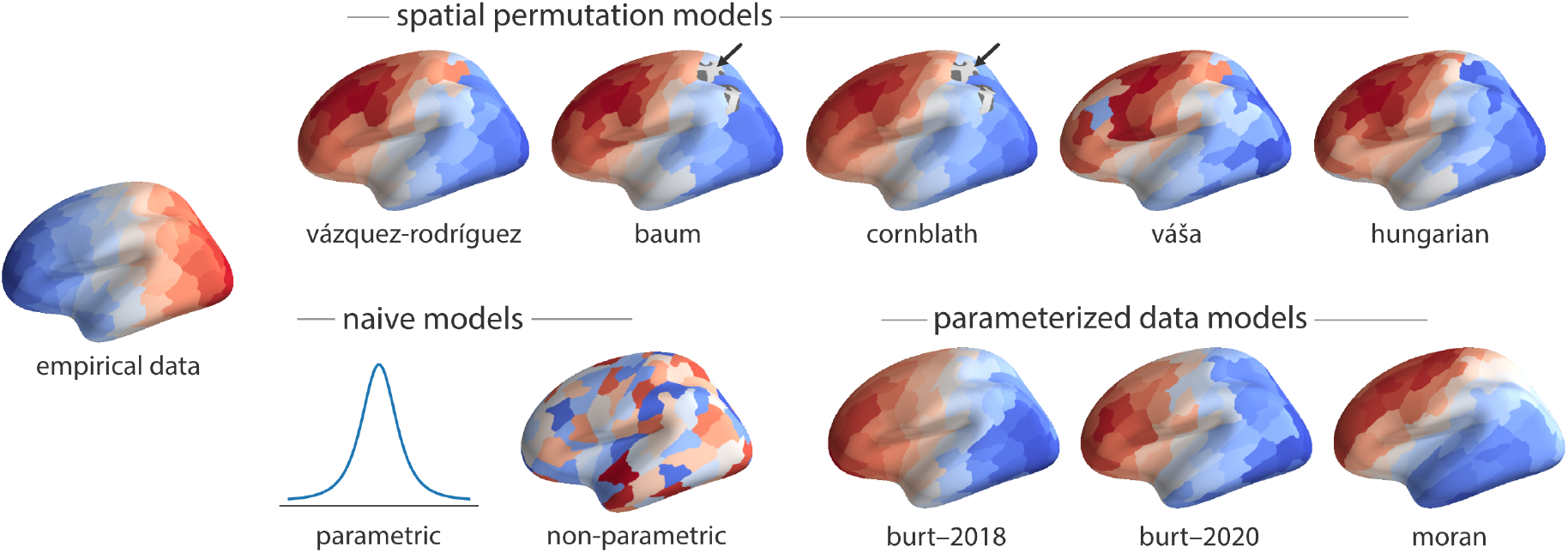
Null model framework examples. Examples of the null frameworks applied to toy data representing an anterior-posterior gradient (empirical data, left) using the 219-parcel Cammoun atlas. All spatial permutation null examples were generated using the same rotation matrix to highlight variations between the different implementations. Parameterized null examples were chosen so as to maximize similarity to the spatial permutation nulls for visualization purposes. Black arrows in the *Baum* and *Cornblath* method examples indicate missing values due to rotation of the medial wall into the cortical surface (see *Methods: Null model frameworks* for more information).

**Figure 2.**
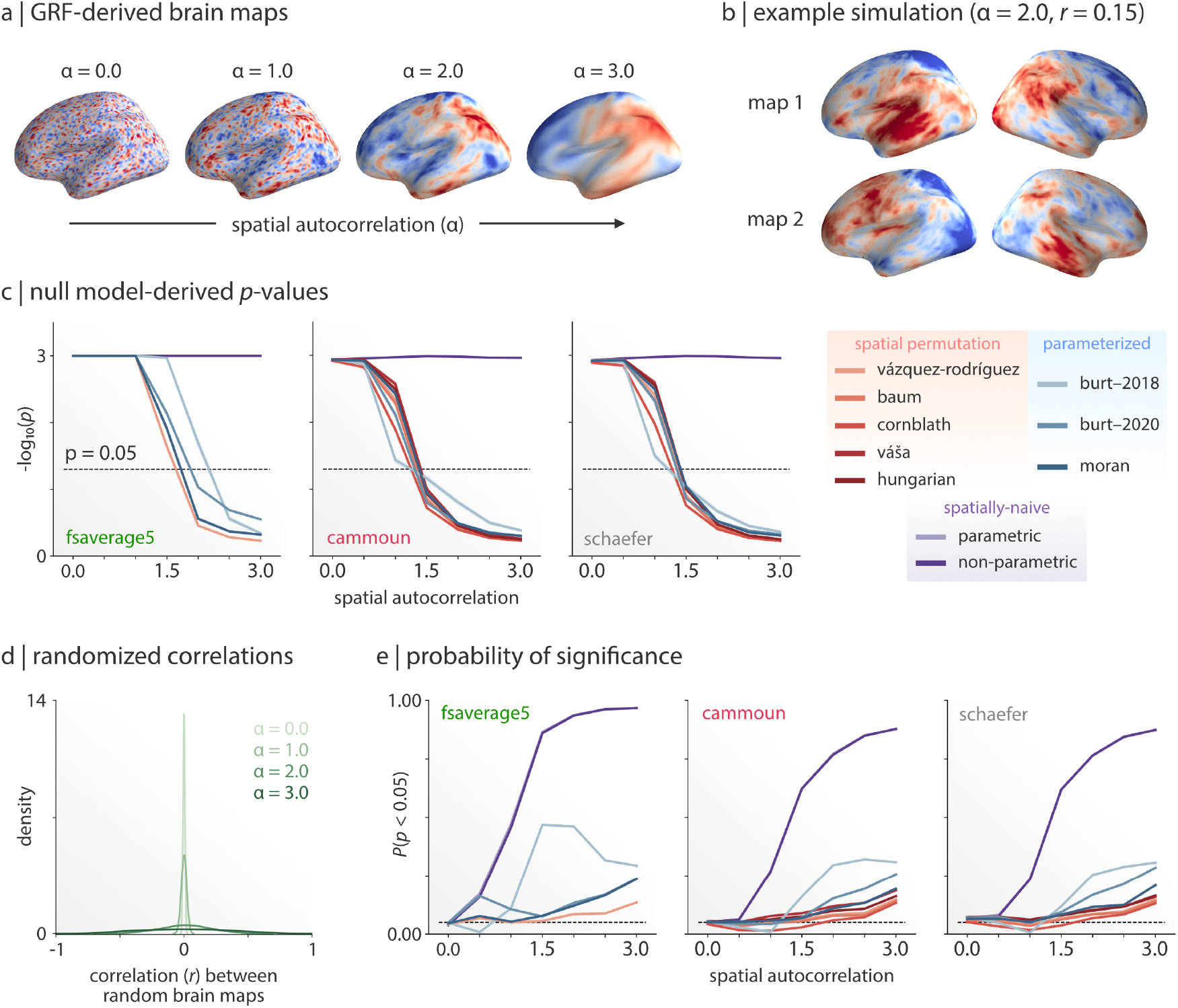
Null model performance on simulated brain maps. (a) Four maps derived from the same Gaussian random field (GRF) with increasing levels of spatial autocorrelation. The map corresponding to *α* = 0.0 represents random Gaussian noise. (b) An example pair of simulated brain maps (*α* = 2.0) correlated at *r* = 0.15 ± 0.005. (c) Average null model performance on 1,000 simulations across all seven levels of spatial autocorrelation. Colored lines and shaded regions on each plot represent the mean and 95% confidence interval. The Cammoun and Schaefer atlas results are shown for the highest resolution (1,000 parcels) only. The dashed black line corresponds to *p* = 0.05 (where values beneath the line indicate *p* > 0.05). The spatially-naive parametric results are not depicted as they are largely indistinguishable from *p* = 0 and approach infinity on the provided scale. Note that the so-called *Vázquez-Rodríguez* method is identical to the framework proposed by Alexander-Bloch et al. (2018) when used at the fsaverage5 resolution; we retain the former name for consistency with the parcellated results, and thus only show vertex-level results for this spatial permutation method. (d) Distributions of correlations between randomized brain map pairs for different levels of spatial autocorrelation. Correlations are shown for vertex (fsaverage5) representations of the data. (e) False positive rate (FPR) for each null model across 1,000 simulations as a function of spatial autocorrelation. Line colors correspond to the legend shown alongside panel (c). The dashed black line corresponds to the expected FPR of five percent. Note that the spatially-naive null models (parametric and non-parametric) are almost completely overlapping on this plot. For the AUC of each null model refer to Table S3.

Examination of *p*-values from the simulated data shows generally comparable performance across spatially-constrained null frameworks (Fig. 2c; red and blue lines). At lower levels of spatial autocorrelation (*α* ≤ 1.5), spatially-constrained null models fare equivalently to spatially-naive models (Fig. 2c; purple lines); however, at higher levels of spatial autocorrelation (*α >* 1.5), spatially-constrained nulls yield more conservative statistics. Although parcellating the vertex-level brain map pairs modifies their correlation structure, parcellated results are consistent with those observed in vertex-wise data (Fig. S4).

Next, to investigate the false-positive rate (FPR) of the null models, we took the simulated brain map pairs from the previous analysis and shuffled them, yielding pairs of randomly-correlated brain maps. At low levels of spatial autocorrelation (*α* ≤ 2), the distribution of correlations between the randomized brain map pairs are tightly centered on *r* = 0.0; however, with increasing levels of spatial autocorrelation (*α >* 2) the variance of the correlation distribution increases dramatically (Fig. 2d).

In other words, at high levels of spatial autocorrelation two randomly-chosen brain maps are more likely to be strongly correlated. To test whether the null frameworks can adequately control for this broadening of the correlation distribution, we re-applied each framework to all randomized pairs of brain maps, generating a *p*-value distribution for each framework. We then assessed the probability with which a given framework would yield a value less than 0.05 (which, in this case, amounts to the false-positive rate; FPR).

When there is no spatial autocorrelation present in the data (*α* = 0.0) all the models perform equivalently, with a FPR around the expected 5%; however, as the spatial autocorrelation increases the FPR of the models begins to diverge (Fig. 2e). Unsurprisingly, spatially-naive models quickly reach a FPR near 100% (Fig. 2e; purple lines), mirroring previously-reported results (Burt et al. 2020). Perhaps more notably: all of the spatially-constrained null models have inflated FPRs at higher levels of spatial autocorrelation (*α* ≥ 2.0; Fig. 2e; red and blue lines). At the highest level of spatial autocorrelation even the most conservative null models have a FPR of approximately 13% (fsaverage5: 13.5%; Cammoun: 13.3%; Schaefer: 13.1%). Moreover, at these higher levels of spatial autocorrelation, significant differences appear between the families of spatially-constrained null models, with the parameterized data models displaying elevated FPR compared to spatial permutation models. We find that the impact of spatial autocorrelation on FPR and these differences between null models is consistent across different parcellation resolutions (Fig. S5). Note also that for the *Burt-2020* model we find that parameter selection seems to influence the resulting FPR (Fig. S2).

To examine whether the observed differences between null model performance arise from poorly-constructed null maps, we examined how well the nulls generated by each framework captured the spatial autocorrelation of the original data. We calculated the Moran’s I of 10,000 simulated brain maps at each level of spatial autocorrelation (Fig. 3a-d, left panel) and, for one simulation at each level of spatial autocorrelation, calculated the Moran’s I of 10,000 null maps generated by each of the null frameworks (Fig. 3a-d, right panels).

**Figure 3.**
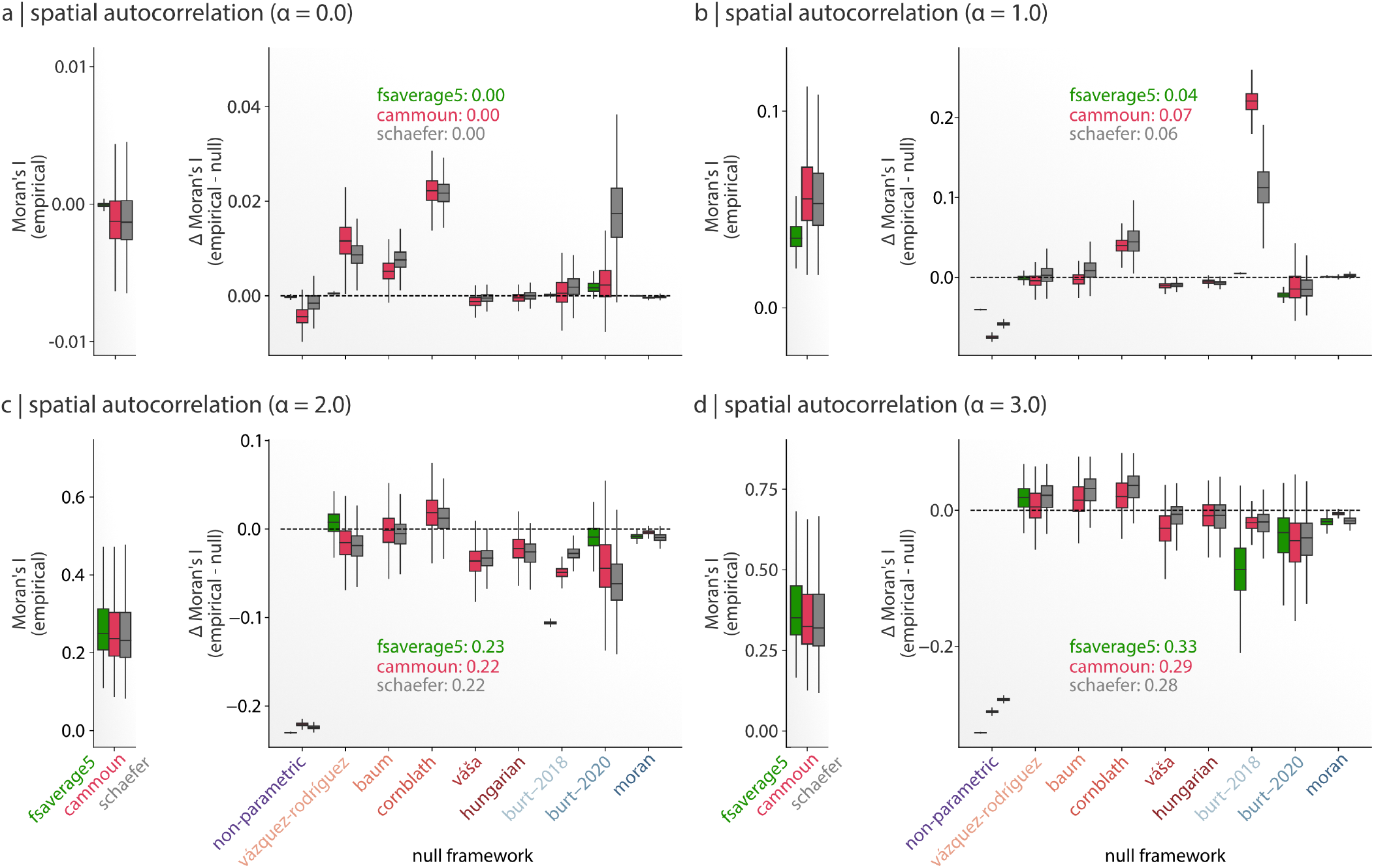
Null models differentially retain autocorrelation of simulated data. Left panel (a-d): spatial autocorrelation estimates of 10,000 simulated maps, estimated via Moran’s I, for fsaverage5, Cammoun atlas (1,000 parcel resolution), and Schaefer atlas (1,000 parcel resolution). Right panel (a-d): spatial autocorrelation estimates of 10,000 null maps, generated from one simulated map, for each null method for all three data resolutions. Colored text in each plot indicates the Moran’s I of the simulated map used to generate the null maps. Plots (a-d) represent data for different levels of spatial autocorrelation of the underlying simulated GRFs (*α* = 0.0, 1.0, 2.0, and 3.0). Note that when applied to vertex-wise fsaverage5 data all of the spatial permutation null methods are identical to the technique proposed in Alexander-Bloch et al. (2018) (see *Methods: Null model frameworks*); we retain the *Vázquez-Rodríguez* label for consistency with results from the parcellated data, and thus only show vertex-level results for this spatial permutation method.

Differences in the Moran’s I of the simulated brain maps are large between the different levels of spatial autocorrelation (*α*)—as would be expected since the *α* is being experimentally modified—but nominal between vertex-wise and parcellated data. The Moran’s I of the null maps, on the other hand, seems to vary as a function of both framework and spatial autocorrelation. Within the spatial permutation null frameworks we observe a general split in method performance: the *Váša* and *Hungarian* methods behave more similarly in comparison to the *Vázquez-Rodríguez*, *Baum*, and *Cornblath* methods. The former methods are performing “perfect” permutations, which come at the expense of the relative spatial orientation (and therefore the spatial autocorrelation) of the input data, whereas the latter methods attempt to more closely retain the spatial orientation at the cost of slightly modifying the underlying data distribution (via e.g., duplicate reassignments). Interestingly, the Moran’s I statistics of the null maps generated with the *Moran* framework—which is designed such that the null maps should *explicitly* match the Moran’s I of the input brain maps—have perhaps the closest correspondence to the input brain maps of all the null models; however, as previously observed, the FPR of the *Moran* method remains categorically higher than the FPR of all the spatial permutation frameworks (Fig. 2e). This discrepancy suggests that optimal signal detection may not solely depend on matching spatial autocorrelation, but also on matching other features of the input brain maps including higher-order correlations or spatial nonstationarities (see *Discussion*).

### Testing correspondence between brain maps

Next, we sought to investigate how the null models perform on two prototypical analyses applied to empirical datasets from the neuroimaging literature. We first examine how each of the ten null models performs when testing the correlations between meta-analytic functional activations. Following Alexander-Bloch et al. (2018), we parcellated meta-analytic “association” maps for 123 cognitive terms derived from NeuroSynth (Poldrack et al. 2011, Yarkoni et al. 2011). We used the term-specific association maps to construct a term × term linear correlation matrix, indicating the extent to which pairs of terms share a similar spatial pattern (Fig. 4a). We then applied each null model to generate a population of null term × term correlation matrices (Fig. 4b). Finally, we constructed a null distribution by retaining the maximum absolute correlation coefficient from each of the null term × term matrices, excluding the diagonal elements (Fig. 4c). This procedure yielded distributions of null correlations for each null model, which were used to threshold the empirical term × term correlation matrix (Alexander-Bloch et al. 2018).

**Figure 4.**
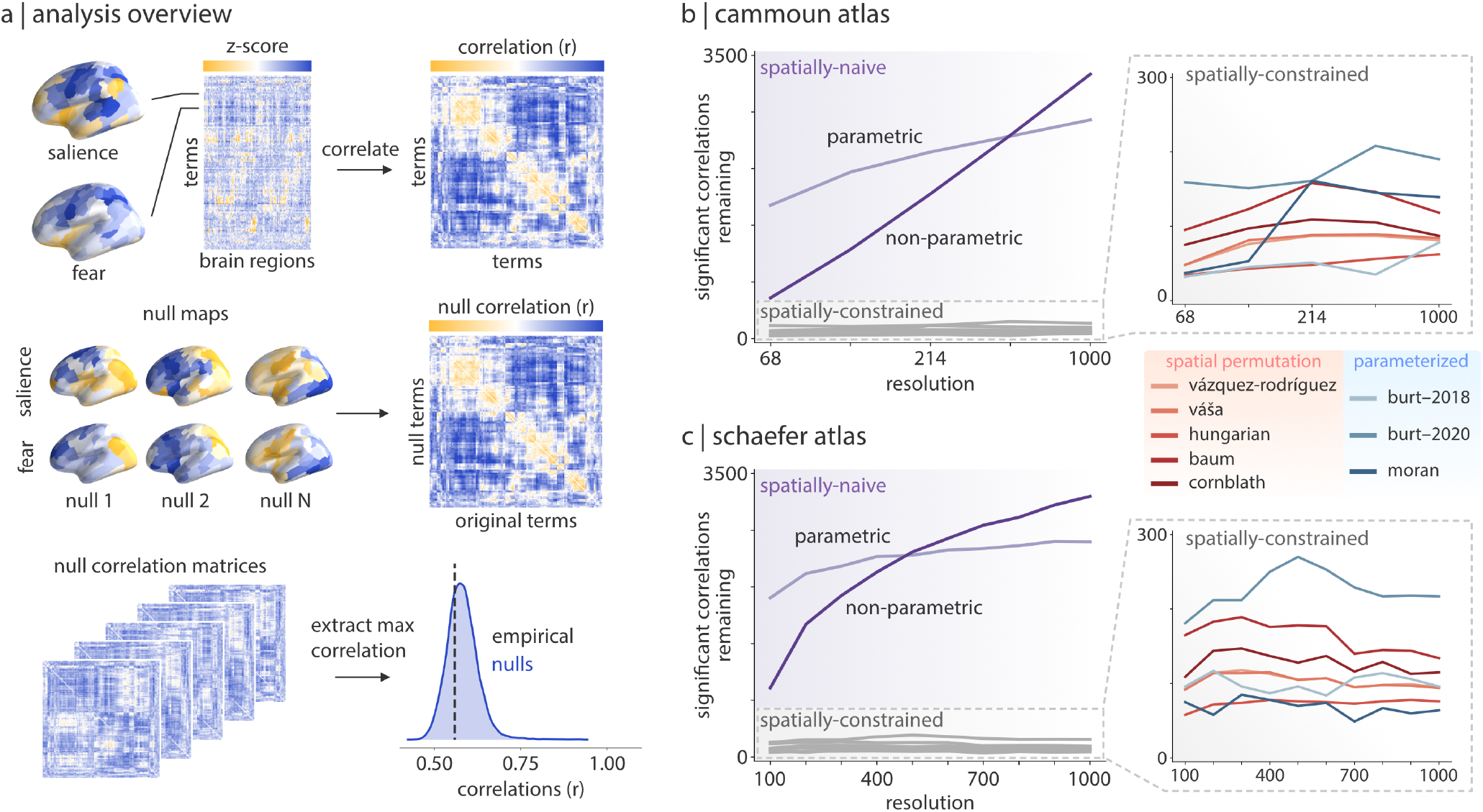
Testing correspondence between brain maps. (a) “Association test” z-score brain maps for 123 terms were downloaded from NeuroSynth. The parcellated term maps were correlated across brain regions to generate a term-by-term correlation matrix. Each null framework was applied to the original term maps to generate 10,000 null maps per term. The original term maps were correlated with each of the null term maps to generate 10,000 null correlation matrices. The maximum absolute correlation in each null matrix (excluding the diagonal) was extracted and retained to create a null distribution of correlations. This null distribution was used to threshold the original correlation matrix at *p* ≤ 0.05. (b, c) The number of significant correlations in the thresholded term matrix for each of the null frameworks as a function of atlas (panel b: Cammoun atlas; panel c: Schaefer atlas) and atlas resolution. Refer to Table S1 for a full list of NeuroSynth terms used.

Fig. 4b,c show the number of statistically significant correlations that remained in the term-by-term matrix after thresholding. All comparisons were performed at *p* ≤ 0.05. To test the interaction between null model and parcellation resolution, we highlight results across all resolutions of the Cammoun and Schaefer parcellations (5 resolutions for the Cammoun atlas; 10 resolutions for the Schaefer atlas).

Altogether, we find notable differences between null models. Mirroring results from our simulation analyses, both the parametric and non-parametric spatially-naive models yield unrealistically liberal results. Within the spatially-constrained nulls, some of the methods are slightly more liberal, yielding higher numbers of significant correlations (e.g. *Burt-2020*, *Váša*), while others are consistently more conservative (*Burt-2018*, *Cornblath*). These differences do not appear to break down according to null model family as they did in the previous analyses, with instances of spatial permutation and parameterized data nulls appearing as both more conservative and more liberal. Moreover, although there are differences among null models, the relative ordering of the null models is stable across multiple parcellation atlases and resolutions, suggesting that models perform consistently across multiple node definitions.

### Testing partition specificity

We next compare null models on tests of partition specificity, in which a researcher examines whether a spatial feature (e.g., cortical thickness, intracortical myelin, functional connectivity strength) is significantly more concentrated in a partition of interest (e.g., in a specific intrinsic functional network or cytoarchitectonic class). Specifically, we tested whether the spatial distribution of the T1w/T2w ratio—thought to reflect intracortical myelin (Glasser and Van Essen 2011)—is circumscribed by intrinsic functional (Yeo et al. 2011) or cytoarchitectonic (Scholtens et al. 2018, von Economo and Koskinas 1925) network boundaries.

We first calculated the mean T1w/T2w ratio for all constituent parcels in the seven intrinsic networks derived by Yeo et al. (2011) (Fig. 5a). We then used each of the null models to generate a distribution of network-specific T1w/T2w means. Finally, each empirical network mean was expressed as a z-score relative to its respective null distribution. A network-specific *p*-value was estimated by computing the proportion of absolute-valued null network means that were greater than the absolute-valued empirical network mean (Fig. 5a), quantifying the probability that the T1w/T2w ratio is significantly greater or smaller in a particular network, above and beyond the network’s size, symmetry, etc.

**Figure 5.**
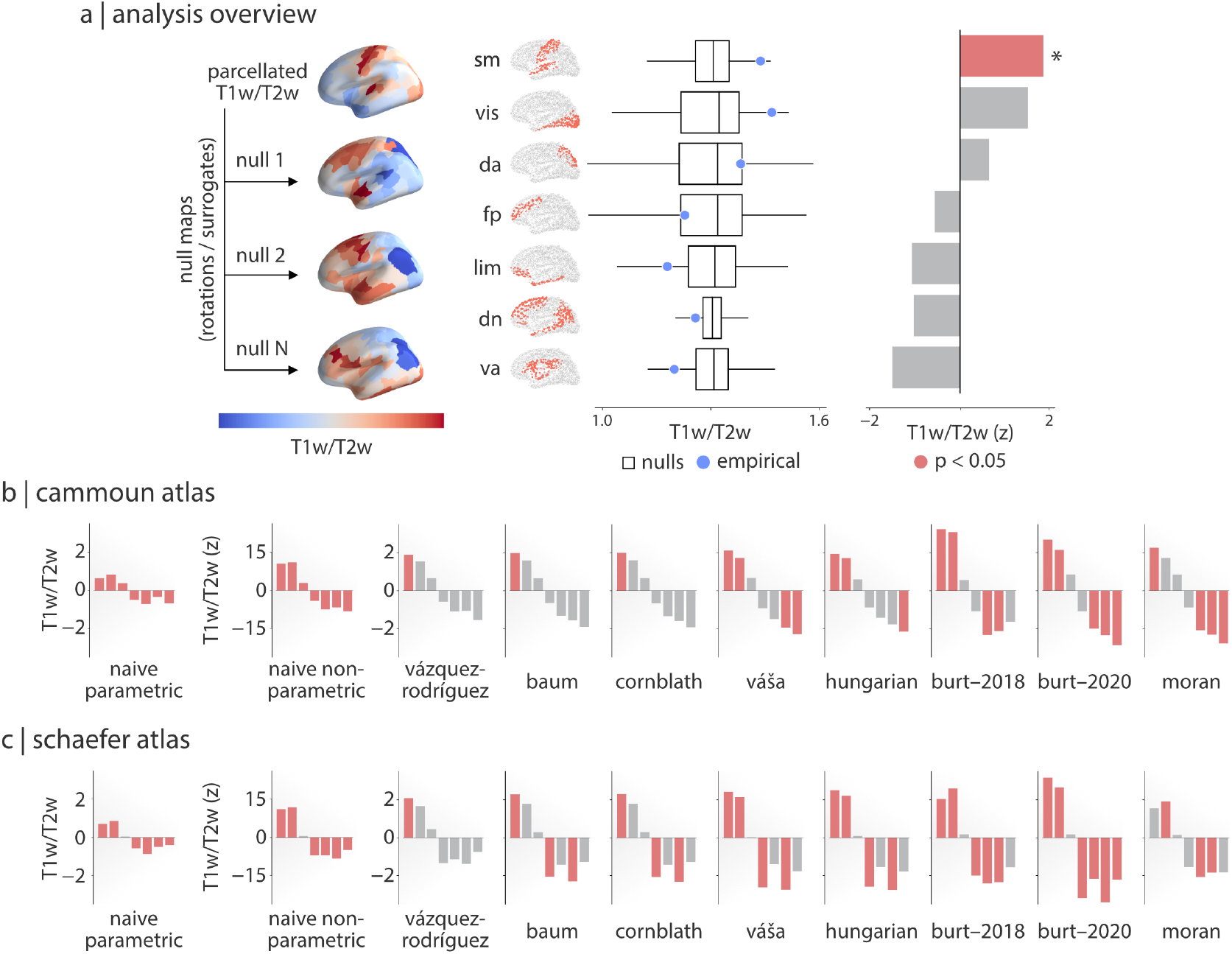
Testing partition specificity. (a) Parcellated T1w/T2w data from the Human Connectome Project were subjected to each of the described null frameworks, yielding 10,000 null T1w/T2w brain maps per framework. Parcels were averaged within each of the seven Yeo-Krienen functional resting-state networks (Yeo et al. 2011) for both the empirical (i.e., real) T1w/T2w brain map (blue dot) and the 10,000 null maps for each framework (white boxplot). Null distributions were used to normalize the empirical values, yielding a single z-score per network. A *p*-value was computed for each network as the proportion of null T1w/T2w network values more extreme than the real T1w/T2w value, divided by the total number of nulls. Networks with a *p*-value < 0.05 are shown in red. (b, c) The procedure described in panel (a) was performed for each of the null frameworks for both the Cammoun (panel b) and Schaefer (panel c) atlases. Results are shown for the highest scale parcellation (1,000 parcels) of each atlas. Yeo-Krienen networks are shown from left-to-right in the same top-to-bottom order as in panel (a). Network assignments: sm = somatomotor, vis = visual, da = dorsal attention, fp = frontoparietal, lim = limbic, dn = default network, va = ventral attention.

Fig. 5b,c shows network-specific z-scores for T1w/T2w ratios. Intrinsic networks with statistically significant (two-tailed, *p* ≤ 0.05) mean T1w/T2w ratios are shown in red, and non-significant networks are shown in grey. We repeated these comparisons for all resolutions of the Cammoun and Schaefer atlases.

We observe three broad trends. First, null models are generally consistent in terms of effect direction and relative ordering of network z-scores: most methods identify over-expression of T1w/T2w ratio in the somatomotor and visual networks, and under-expression of T1w/T2w ratios in the ventral attention, default, frontoparietal and limbic networks (Fig. 5b,c). Second, despite these similarities, individual null models are inconsistent in the statistical inferences they provide: the number and identity of significant networks varies considerably from model to model. Third, the models are variable in terms of how conservative they are. Mirroring results from simulations, and in contrast to results using NeuroSynth data, differences among models appear to largely break down according to model family, with spatial permutation null models tending to be more conservative (*Vázquez-Rodríguez*, *Baum*, *Cornblath*) and parameterized nulls more liberal (e.g., *Burt-2018* and *Burt-2020*). These distinctions between model families are additionally apparent in the shape of the underlying null distributions generated by each framework (Fig. S7).

We also investigate differences between parcellation resolution and network partitions. Fig. 6a shows results for a structural and functional atlas (Cammoun and Schaefer) across 5 and 10 resolutions, respectively. The means are expressed as z-scores relative to a null distribution generated by a given method, and were computed for seven intrinsic functional networks (Yeo et al. 2011), as well as seven cytoarchitectonic classes (Scholtens et al. 2018, von Economo and Koskinas 1925) (Fig. 6b). The null model-, parcellation-, resolution- and partition-specific z-scores are displayed in Fig. 6c and d. Overall, the findings are consistent with the intuitions drawn above. Namely, we continue to observe consistency across nulls in terms of effect direction and ordering, and inconsistency in the number and identity of significant partitions; however, there is less differentiation between the spatial permutation and parameterized data null model families for the von Economo partition classes. This consistency across null frameworks may indicate that the von Economo partitions provide a truer or more accurate delineation of T1w/T2w data than the Yeo-Krienen networks. Although not specifically related to differences between models, we also note an example of how atlas and partition may potentially interact and lead to different inferences: class #7 of the von Economo partition (limbic regions) is consistently deemed to be significantly under-enriched for T1w/T2w across Cammoun-derived parcellations, while it is generally not statistically significant across Schaefer-derived parcellations (Fig. 6d).

**Figure 6.**
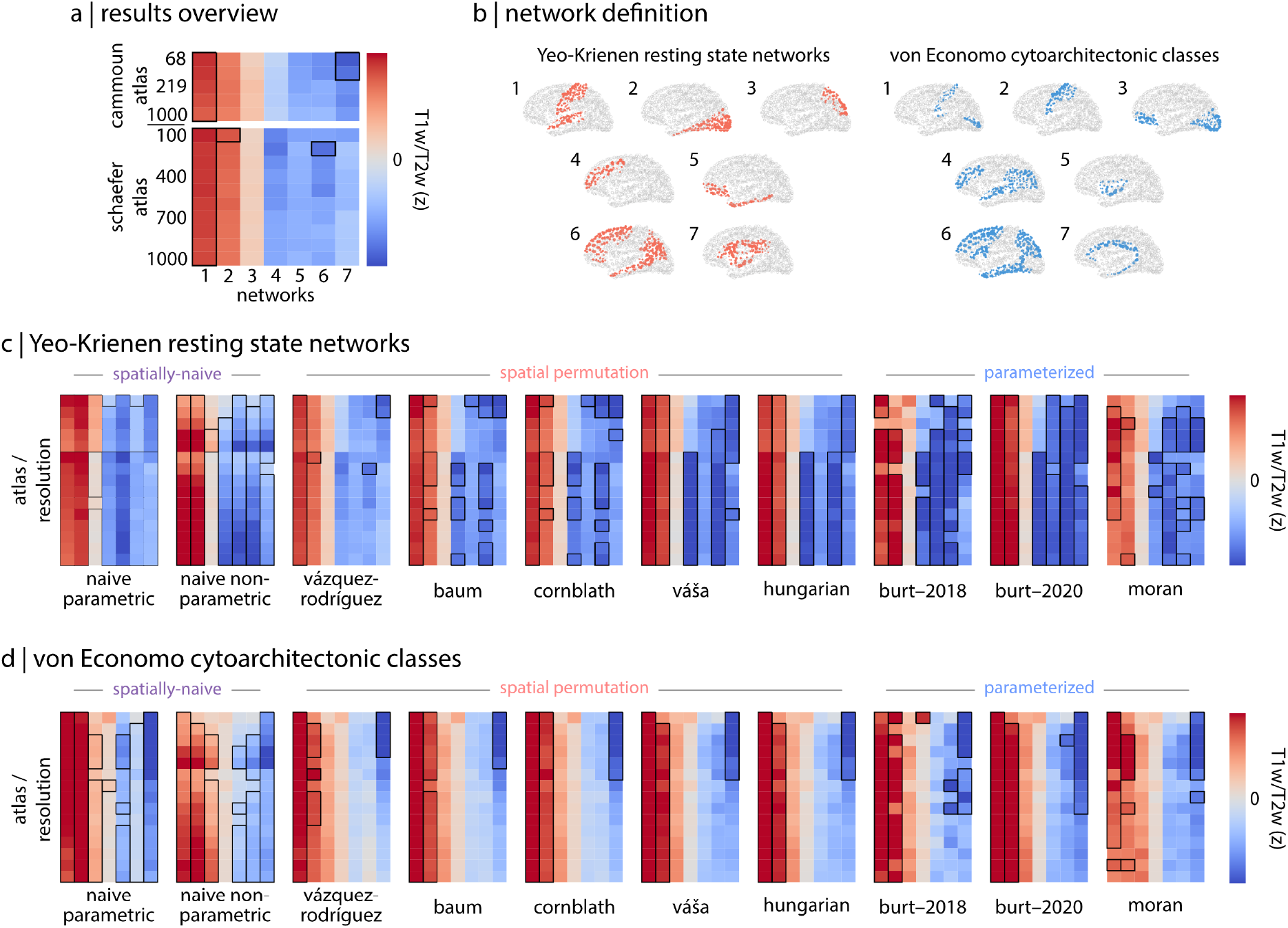
Partition-specific definition of T1w/T2s maps. (a) Overview of the results assessing partition specificity of T1w/T2w measurements. Each cell in the heatmap represents the T1w/T2w z-score for a given network (x-axis) and atlas / resolution (y-axis). Black outlines around a cell represent network z-scores that are significant at the *p* ≤ 0.05 level. Note that the fifth and fifteenth row of each heatmap represent the same values shown in Fig. 5b. (b) The network labels for both the seven Yeo-Krienen resting state networks (Yeo et al. 2011) and the seven von Economo-Koskinas cytoarchitectonic classes (Scholtens et al. 2018, von Economo and Koskinas 1925). (c, d) The results for all ten null models for all atlases and resolutions for the (c) Yeo-Krienen resting state networks and (d) von Economo cytoarchitectonic classes. While there is some notable variation amongst atlases and resolutions, primary differences are observable between null frameworks and partitions. Naive parametric heatmaps represent partition means, not z-scores, and values range from −1–+1. Naive non-parametric heatmap values range from −7.5–+7.5, while all other null model heatmaps range from −2.5–+2.5]. Yeo-Krienen resting state networks: 1 = somatomotor, 2 = visual, 3 = dorsal attention, 4 = frontoparietal, 5 = limbic, 6 = default, 7 = ventral attention. von Economo cytoarchitectonic classes: 1 = primary sensory cortex, 2 = primary motor cortex, 3 = primary/secondary sensory cortex, 4 = association cortex, 5 = insular cortex, 6 = association cortex, 7 = limbic regions.

One notable disadvantage of the *Baum* and *Cornblath* spatial permutation null models is that the rotations on which they rely will cause the medial wall to be displaced into the cortical sheet. The resulting null distributions can have missing data for this portion of the cortex (Fig. 1), yielding uneven sample sizes between null distributions and potentially biasing results, especially if one partition or networks is impacted more than others. To investigate whether this may have influenced the presented results we re-ran all partition specificity analyses for the *Baum* and *Cornblath* methods, discarding rotations where there was excessive data loss due to the rotation of the medial wall, and find comparable results (Fig. S6).

### Implementation of null models can impact performance

While the presented results suggest that, at a broad level, selection of null model framework is an important choice for researchers, there are additional choices to be made within each framework. Here, we investigate how different implementations and seemingly minor methodological choices of spatial permutation and parameterized data null models can influence statistical outcomes.

#### Variability in parcel centroid definition

Of the five spatial permutation null models presented in the current report, three require definition of a centroid for each parcel (*Vázquez-Rodríguez*, *Váša*, and *Hungarian*). For the analyses reported, parcel centroids were generated following the procedure used by Vázquez-Rodríguez et al. (2019), which include: (1) taking the average of the coordinates of all vertices within a given parcel and (2) using the coordinates of the surface vertex closest to this average (where closest is defined as minimizing Euclidean distance). However, Váša et al. (2018) defined their parcel centroids using only the average of the vertices within each parcel (i.e., not projecting back to a surface vertex). Notably, both of these procedures fail to take into account oblong or C-shaped parcels for which the center-of-mass may fall outside the boundaries of the parcel. An alternative approach to account for this possibility is to find the coordinates of the vertex that minimizes the average surface (geodesic) distance to all other vertices within each parcel.

We assessed the extent to which these three different methods of parcel centroid definition impacted the generated rotations for the three relevant null models. We generated ten rotation matrices and applied them to the coordinates derived from each of these three parcel centroid definition methods, reassigning parcels using the approach from each of the three relevant null frameworks. We compared the similarity of the reassignments using the normalized Hamming distance (Fig. 7a; Hamming 1950). Critically, because the same rotations were applied for every model and method, any observed differences are a result of variation in either the parcel centroid coordinates or models themselves.

**Figure 7.**
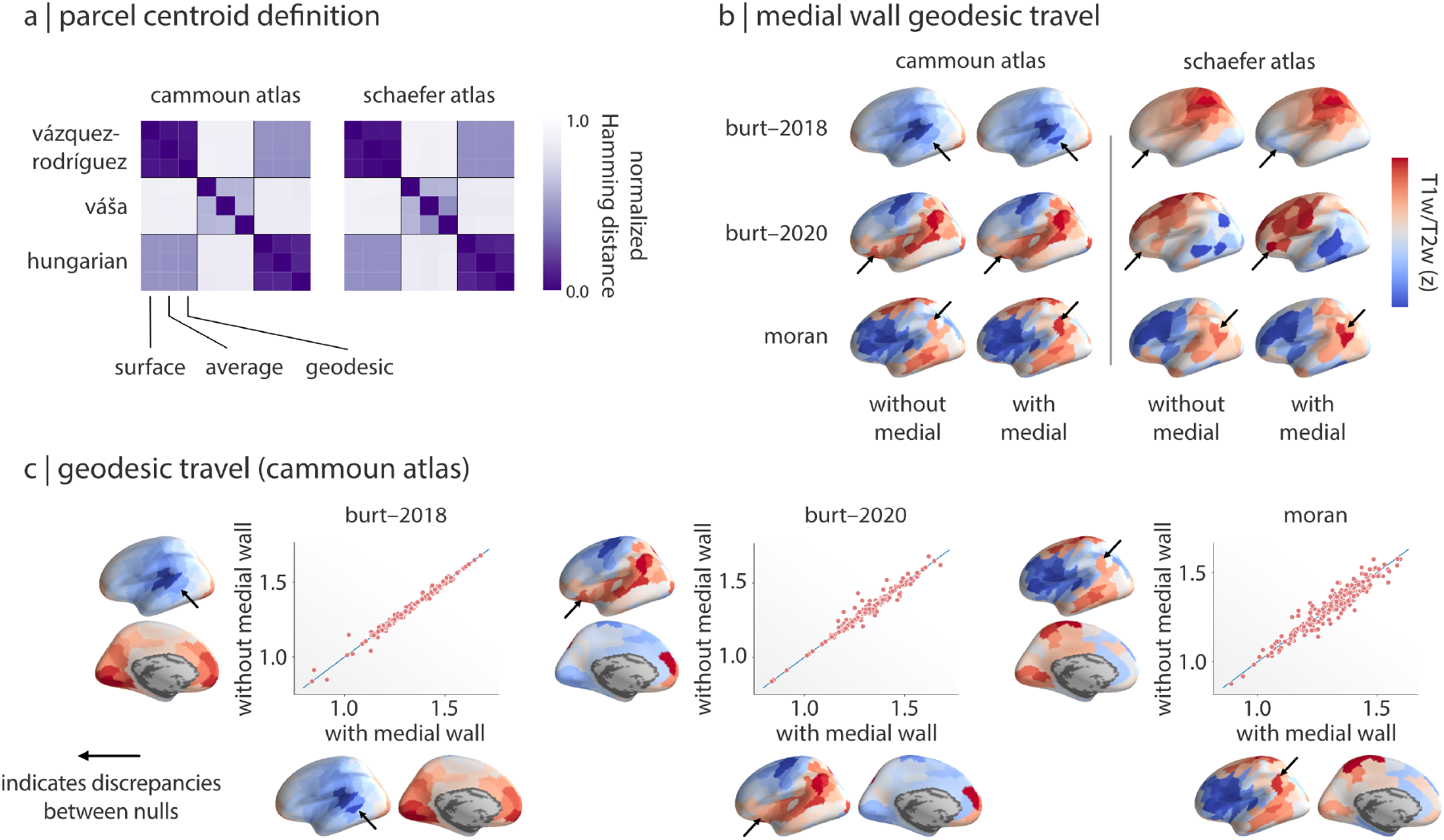
Implementation of null models can impact performance. (a) The impact of parcel centroid definition on three spatial permutation null models. Heatmaps show the correspondence (normalized Hamming distance) between null reassignments generated using different null frameworks and parcel centroid methods, where lower values (purple) indicate greater correspondence. The heatmaps are broken into nine sections (black outlines) which delineate different null frameworks; cells within each section represent the different parcel centroid methods. For results from all resolutions of the Cammoun and Schaefer atlases refer to Fig. S8. (b) Example surrogates generated from distances matrices allowing (“with medial”) and disallowing (“without medial”) travel along the medial wall for the three parameterized data null models (*Burt–2018*, *Burt–2020*, and *Moran*). Arrows indicate parcels where values are discrepant between pairs of surrogates for each method. (c) Scatterplots of parcel values for surrogates generated with / without medial wall travel. Note: “Cammoun atlas” refers to the 219-parcel resolution and “Schaefer atlas” to the 200-parcel resolution.

Fig. 7a shows the Hamming distance between reassignments for all combinations of these parcel centroid and null framework methods across the Cammoun and Schaefer atlases (219 and 200 nodes, respectively; refer to Fig. S8 for all resolutions). Predictably, we observe the strongest differences between reassignments generated from the different null methods. In particular, the *Váša* null method seems to markedly differ from the *Vázquez-Rodríguez* and *Hungarian* methods. In addition, there is also variation between parcel centroid calculation methods; for example, the *Váša* method seems to have only moderate correspondence between different parcel centroid definitions. These results highlight important differences not only between spatial permutation null models but within the implementation of each model itself.

#### Geodesic distances along the medial wall

Parameterized data models rely on a user-defined weight matrix that provides an estimation of the spatial structure of the related brain maps. In most cases—as in the current report—this weight matrix takes the form of a geodesic- or surface-based distance matrix calculated from shortest paths taken along the surface mesh projection of a brain. However, the calculation of these distance matrices often fails to take into account that paths along the brain’s medial wall, where the cortical sheet is discontinuous due to underlying white matter and sub-cortical nuclei, should not be possible. Failing to account for this discontinuity yields distance matrices that categorically underestimate the actual distance between many brain regions.

We tested the extent to which generating distance matrices (dis-)allowing travel along the medial wall biases the surrogate data maps generated from the three parameterized null models. Fig. 7b shows an example surrogate map for these null models for the Cammoun and Schaefer atlases. Overall, surrogates generated from the different distance matrices show strong similarities. To examine this in more detail we correlated 1,000 surrogates generated from the two distance matrices for each method, finding remarkably high, though not perfect, correspondence: *r* = 0.9618 [0.9615–0.9622] (*Burt-2018*), 0.9379 [0.9373–0.9384] (*Burt-2020*), 0.9481 [0.9479–0.9482] (*Moran*) (mean [2.5–97.5% CI] across all atlases and resolutions, derived from inverse *r*-to-*z*-transformed distributions). Although parameterized data nulls seem relatively robust to this issue, researchers should be explicit about how they are handling the medial wall when constructing the required distance matrices in their own analyses.

## DISCUSSION

In the present report we systematically assess the performance of ten null frameworks on statistical inference in four analyses. We first examine their efficacy in the context of two comprehensive simulation analyses, testing the accuracy of the statistical estimates and false positive rates generated by each model. Next, we compare the performance of the null models at replicating previously published analyses of two empirical datasets. We close with an investigation of how implementation choices within each null framework may influence their behavior.

Across both simulated and empirical datasets we find that spatially-naive null models consistently yield inflated error rates, unrealistically liberal statistical estimates, and a strong dependence on parcellation resolution, confirming previous reports that such methods are unsuitable for use in neuroimaging (Alexander-Bloch et al. 2018, Burt et al. 2020). In these same analyses we demonstrate that spatially-constrained null models yield more realistic, conservative statistical estimates. However, we also show that even these frameworks suffer from inflated false positives—with error rates as high as 46% in realistic simulations—and that there is considerable variability in their performance. Altogether, these findings suggest that while spatially-constrained null models are the current state-of-the-art, there are significant improvements to be made in this space. The present report provides a harmonised framework against which researchers can benchmark and compare future developments.

Our analyses highlight a consistent distinction in the performance of the spatially-constrained null models: in general, we find that the spatial permutation models tend to yield more conservative statistical estimates and lower error rates than the parameterized data models. Importantly, this variability seems to depend on research context. Although analyses of simulated data show a reliable and marked difference between model families (Fig. 2), this delineation becomes less consistent when examining empirical results. That is, we observe lower variability between nulls when examining brain map correspondences in the NeuroSynth dataset (Fig. 4) than when exploring partition specificity in the HCP dataset (Figs. 5 and 6). Moreover, unlike results from both the simulation and HCP analyses, differences in null model behavior in the NeuroSynth analysis do not break down along the lines of model family.

What is driving these differences between null model families? One possible explanation is the degree of spatial autocorrelation. Results from our simulation analyses (Figs. 2 and 3) highlight that the amount or degree of spatial autocorrelation present in the data has a strong influence on how the null models perform: at lower levels the frameworks all perform quite comparably, with differences emerging only at higher levels. Although this does not explain the variability between model families in-and-of itself, differences in empirical spatial autocorrelation may be partially responsible for the seemingly contradictory results in the NeuroSynth and HCP datasets.

Another possibility is that the null model families embody different hypotheses. On the one hand, spatial permutation models assume that the spatial orientation of two brain maps is responsible for any observed relationships between them, and test this by altering the alignment of the brain maps via random angular rotations. The parameterized data models instead operate under the hypothesis that spatial autocorrelation is driving observed relationships between brain maps, and so test this by generating null maps that aim to match the degree of spatial autocorrelation present in the original data. Although this is a subtle difference—spatial orientation versus spatial autocorrelation—it can have notable consequences. For example, we find that null maps generated by spatial permutation models tend to more accurately match the spatial autocorrelation of the original data than those created with the parameterized data models, despite not explicitly optimizing this factor (Fig. 3). This difference may be especially important in explaining variability of model performance in the presence of spatial non-stationarities commonly found in empirical data.

The assumption of stationarity underlies all of the spatial null models examined. That is, they assume that any spatial autocorrelation present in the data is consistent and uniform throughout; in a system as complex as the brain, this is unlikely to be true. However, although spatial stationarity is an explicit and strict assumption of the parameterized data models, it is a much looser assumption for the spatial permutation models. These rotation-based models should, in theory, retain non-stationarities present in the original data, perhaps yielding more realistic null distributions (though this will vary based on the exact method employed and whether data are discarded due to the medial wall). Future work examining and developing spatial null models should aim to explicitly address the concept of spatial non-stationarities and examine how their presence might impact model performance. Note that one existing method—wavelet-based phase randomization (Breakspear et al. 2003, 2004)— may be well-suited to address this; however, this method has not yet been extended to surface data and so we were unable to examine it in the current report.

Beyond the impact of spatial autocorrelation and spatial non-statitionarities, implementation differences within null families entail additional methodological choices that may explain additional variability in the observed results. Within the spatial permutation nulls, implementation differences are primarily driven by whether the methods are performing a “perfect” permutation of the data (i.e., *Váša*, *Hungarian*) or using a more flexible re-assignment (i.e., *Vázquez-Rodríguez*, *Baum*, *Cornblath*). Using the so-called perfect permutation methods comes at the expense of the relative spatial orientation of the input data and tends to yield slightly more liberal statistical estimates (Figs. 2 and 6). The parameterized nulls mostly vary in how they implement the data-generating process, including their assumptions about the nature of spatial autocorrelation and how it is estimated in the original data. Our results clearly show that these implementation differences have an observable effect on generated statistical estimates, but more research is needed to better understand how differences in performance arise from specific methodological choices.

Encouragingly, our results show negligible differences in spatially-constrained null model performance between “dense” (i.e., vertex-wise) and parcellated data, and furthermore suggest a minimal impact of parcellation and parcellation resolution (Figs. 2, 3, and S5). How brain regions are defined and ultimately represented in analyses is an important choice in neuroimaging (Eickhoff et al. 2018), as recent work has highlighted the role of brain region definition in analyses including test-retest reliability (Arslan et al. 2018), structure-function coupling (Messé 2020, Vázquez-Rodríguez et al. 2019), individual fingerprinting (Finn et al. 2015), modelling of brain dynamics (Proix et al. 2016), and prediction of behavior and disease (Dadi et al. 2020, Kong et al. 2019). Although choice of parcellation is clearly an important methodological consideration, our results suggest that it does not strongly influence spatially-constrained null models. This “parcellation invariance” will hopefully support broader adoption of these null models in future studies.

More generally, this investigation builds on increasing efforts in the neuroimaging literature to benchmark the effects of methodological choices. Recently, the broad impacts of these choices has been demonstrated in research using structural MRI (Bhagwat et al. 2020, Kharabian Masouleh et al. 2020), task fMRI (Botvinik-Nezer et al. 2020, Carp 2012), resting state fMRI (Ciric et al. 2017, Parkes et al. 2018), and diffusion MRI (Maier-Hein et al. 2017, Oldham et al. 2020, Schilling et al. 2019). Concomitant with an increasing awareness of the importance of these choices is a developing trend to share and analyze “raw” or un-thresholded brain maps (Gorgolewski et al. 2015, Witt et al. 2020). Convergence of these trends has naturally opened new research questions that revolve around comparing such brain maps. Methods that appropriately consider the inherent spatial structure of the brain are thus going to be critical to ensuring valid inferences can be drawn from these lines of inquiry. The present study is a first step towards better understanding the variable implementation of these statistical methods and ultimately standardizing or synthesizing their application.

Finally, to help guide researchers in choosing an appropriate null model in their own work, we end with a set of nine important considerations:

1. Spatially-naive null models—both parametric and non-parametric—are inappropriate for significance testing of neuroimaging brain maps. When applied to spatially-autocorrelated brain maps typical of most neuroimaging data these models approach a false positive rate of >75%. Use of spatially-naive null models should be actively discouraged throughout the field.
2. Parameterized data models are necessary when working with voxel-wise, subcortical, cerebellar, and region-of-interest data, as spatial permutation null models are only viable for data that can be projected to a spherical representation of the cortical surface. Note that within the parameterized data models some may be less feasible for use with high-resolution datasets (e.g., the Moran method relies on an eigendecomposition, which can be computationally prohibitive in some cases).
3. Spatial permutation frameworks tend to have lower overall error rates than parameterized data models; however, this difference is less pronounced for data with low levels of spatial autocorrelation. Appropriate choice of null framework will thus depend on research context.
4. Parcellation and parcellation resolution appear to have negligible impact on the performance of spatially-constrained null models and limited impact on statistical inference. Researchers should use whatever parcellation and resolution best suits their data and research questions.
5. Parameterized data models have a greater computational cost than spatial permutation models (Fig. S1). When working with parcellated data this difference is less pronounced and the computational cost of the two families becomes more comparable (though does not equalize).
6. Spatial permutation null models are only viable for data that can be projected to a spherical representation of the cortical surface. This precludes use of these null models with voxel-wise (unless a volume-to-surface projection is used), subcortical, cerebellar, and region-of-interest data; in these cases parameterized data models should be used. Note that within the parameterized data models some may be less feasible for use with ultra high-resolution datasets (e.g., the *Moran* method relies on an eigendecomposition, which can be computationally prohibitive in some cases.)
7. Avoid relying on default hyperparameter choices when using parameterized data models (i.e., *Burt-2020*). Selecting parameters to ensure that the null models are providing a good fit to the data can reduce false positive rates by nearly half (Fig. S2).
8. Spatial null models can be applied to many analysis procedures, including ones that are not often statistically assessed—such as the partition specificity analysis presented in this article. Simply reporting the average values of a given feature within a set of partitions can misrepresent which partitions are truly over- or under-expressing the feature. We find that in the presence of spatial autocorrelation these descriptive estimates can be misleading, and so recommend using spatial null models to statistically assess these values.
9. When retaining the exact distribution of the original brain map is important, such as when testing network-based statistics like degree, the *Váša* or *Hungarian* spatial permutation models should preferentially be used. These models attempt to perfectly preserve the distribution of the original data and are more well-suited to these types of hypotheses.
10. Null models converge quickly (i.e., with relatively few permutations / rotations / surrogates; Fig. S3), reaching stable statistical estimates after approximately 100–500 nulls. Researchers can use this information to balance accuracy and computational feasibility when selecting the number of nulls to use in their analysis.

## CONCLUSIONS

The present report comprehensively examines ten null models for comparing brain maps in neuroimaging research. We find dramatically improved performance and reduced error rates in null models that account for spatial autocorrelation when compared with spatially-naive models; however, we note that potentially meaningful differences between spatially-constrained nulls are present across all analyses. Our results highlight the need to closely consider the variability across implementation of these null methods and to more explicitly compare their performance across a wider range of research contexts.

## ACKNOWLEDGEMENTS

We thank Laura Suárez, Golia Shafiei, Justine Hansen, Vincent Bazinet, Bertha Vázquez-Rodríguez, Elizabeth DuPre, and Joshua Burt for their comments and suggestions. This research was undertaken thanks in part to funding from the Canada First Research Excellence Fund, awarded to McGill University for the Healthy Brains for Healthy Lives initiative. This work was supported in part by funding provided by Brain Canada, in partnership with Health Canada, for the Canadian Open Neuroscience Platform initiative. RDM acknowledges support from the Fonds du Recherche Québec - Nature et Technologies and the Canadian Open Neuroscience Platform. BM acknowledges support from the Natural Sciences and Engineering Research Council of Canada (NSERC Discovery Grant RGPIN #017-04265) and from the Canada Research Chairs Program.

---

**Table S1.** List of terms used in NeuroSynth analyses. Terms that overlapped between NeuroSynth (Yarkoni et al. 2011) and Cognitive Atlas (Poldrack et al. 2011) corpuses used in the reported analyses. For a machine-readable format please refer to https://github.com/netneurolab/markello_spatialnulls.

|  |  |  |  |  |
| --- | --- | --- | --- | --- |
| action | eating | insight | naming | semantic memory |
| adaptation | efficiency | integration | navigation | sentence comprehension |
| addiction | effort | intelligence | object recognition | skill |
| anticipation | emotion | intention | pain | sleep |
| anxiety | emotion regulation | interference | perception | social cognition |
| arousal | empathy | judgment | planning | spatial attention |
| association | encoding | knowledge | priming | speech perception |
| attention | episodic memory | language | psychosis | speech production |
| autobiographical memory | expectancy | language comprehension | reading | strategy |
| balance | expertise | learning | reasoning | strength |
| belief | extinction | listening | recall | stress |
| categorization | face recognition | localization | recognition | sustained attention |
| cognitive control | facial expression | loss | rehearsal | task difficulty |
| communication | familiarity | maintenance | reinforcement learning | thought |
| competition | fear | manipulation | response inhibition | uncertainty |
| concept | fixation | meaning | response selection | updating |
| consciousness | focus | memory | retention | utility |
| consolidation | gaze | memory retrieval | retrieval | valence |
| context | goal | mental imagery | reward anticipation | verbal fluency |
| coordination | hyperactivity | monitoring | rhythm | visual attention |
| decision | imagery | mood | risk | visual perception |
| decision making | impulsivity | morphology | rule | word recognition |
| detection | induction | motor control | salience | working memory |
| discrimination | inference | movement | search |  |
| distraction | inhibition | multisensory | selective attention |  |

**Table S2.** Null framework limitations. Overview of some of the drawbacks of each of the null frameworks described in the current report. Each of the families also suffer from drawbacks impacting all constituent null frameworks (i.e., spatial permutation models can only handle surface data, parameterized models tend to be more computationally complex); refer to *Discussion* for more information.

| Naive models |  |
| --- | --- |
| *Method* | *Drawbacks* |
| Parametric | Assumes errors are *i.i.d.*; does not account for spatial structure |
| Non-parametric | Does not account for spatial structure |
| Spatial permutation models |  |
| *Method* | *Drawbacks* |
| Vázquez-Rodríguez | Allows duplicate parcel assignments |
| Baum | Projection to vertices involves upsampling; allows duplicate parcel assignments |
| Cornblath | Projection to vertices involves upsampling; null data will not match original due to re-averaging |
| Váša | Generates sub-optimal assignments with respect to global cost |
| Hungarian | Undesirable, high-cost (i.e., distant) assignments can be made to optimize global cost |
| Parameterized data models |  |
| *Method* | *Drawbacks* |
| Burt-2018 | Assumes exponential distance model provides accurate estimation of $\hat{\rho}$ and $\hat{d}_0$ from data |
| Burt-2020 | Variogram models may not properly model some spatial distributions |
| Moran | Dependent on provided weight matrix, $\mathbf{W}$, which varies based on neighborhood size; tends to yield highly auto-correlated surrogates |

**Table S3.** AUC of null model false positive rates. The AUC (area under the curve) for the false positive rates of each null model as a function of spatial autocorrelation. Values were calculated using the trapezoidal rule with data presented in Fig. 2e and are normalized such that a false positive rate of 100% across all levels of spatial autocorrelation would be equal to 1. Lower values indicate more nominal error rates; the lowest value in each column is highlighted in bold. Note that the so-called *Vázquez-Rodríguez* method is identical to the framework proposed by Alexander-Bloch et al. (2018) when used at the fsaverage5 resolution; we retain the former name for consistency with the parcellated results.

|  | **fsaverage5** | **Cammoun** | **Schaefer** |
| --- | --- | --- | --- |
| Naive parametric | 0.653 | 0.507 | 0.503 |
| Naive non-parametric | 0.646 | 0.506 | 0.502 |
| Vázquez-Rodríguez | **0.073** | 0.071 | 0.078 |
| Baum | — | 0.065 | 0.071 |
| Cornblath | — | **0.046** | **0.055** |
| Váša | — | 0.099 | 0.092 |
| Hungarian | — | 0.084 | 0.090 |
| Burt-2018 | 0.256 | 0.166 | 0.151 |
| Burt-2020 | 0.131 | 0.110 | 0.129 |
| Moran | 0.106 | 0.090 | 0.096 |

**Figure S1.**
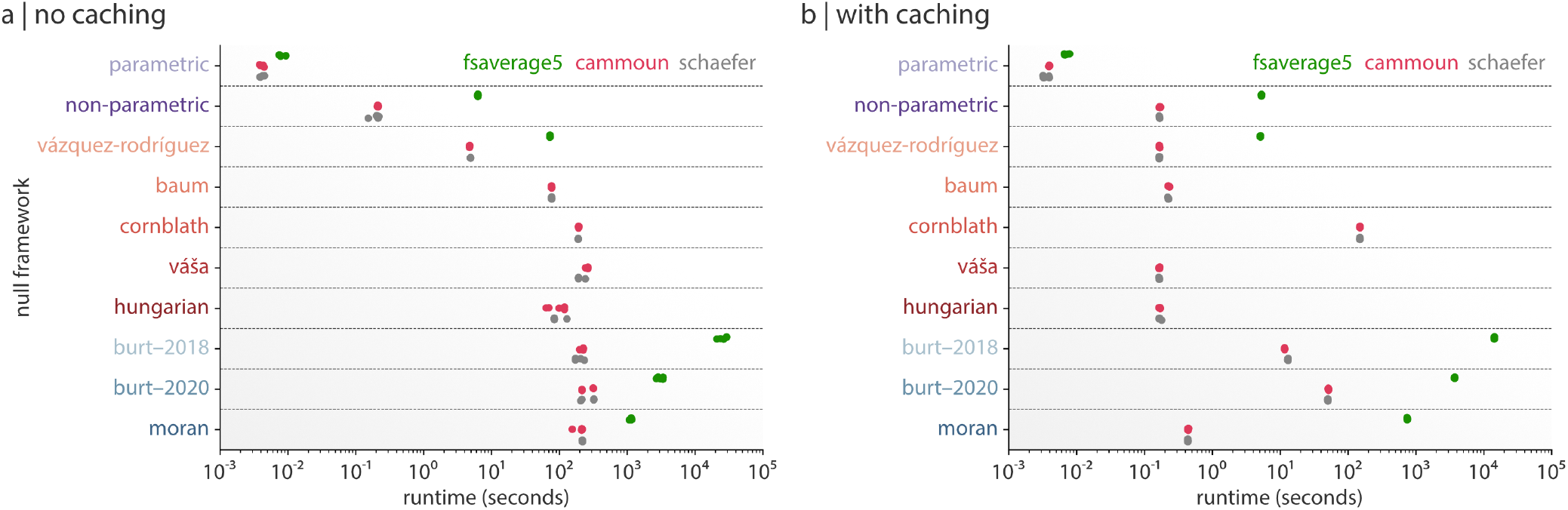
Computational runtime of null frameworks. The computation time of each null framework for the fsaverage5-resolution data (green) and 1,000-parcel Cammoun (pink) and Schaefer (grey) atlases. Each null framework was run five times on a single simulated brain map (*α* = 2.0) and used to generate 1,000 null maps per run; repeats are plotted as separate dots. (a) The computation times of the null frameworks when run with “no caching”, which includes the time required to both generate 1,000 null maps and any relevant intermediate data, such as permutations (non-parametric), spatial rotations (*Vázquez-Rodríguez*, *Váša*, *Hungarian*, *Baum*, *Cornblath*), and geodesic distance matrices (*Burt-2018*, *Burt-2020*, *Moran*), and are limited to one CPU / one thread. (b) The computation time of each null framework when using cached versions of intermediate data (i.e., permutations, spatial rotations, and geodesic distance matrices). Note the decrease in computation time from (a) and, e.g., the equalizing of most of the spatial permutation null models. Results are shown for the 1,000-parcel resolution of the Cammoun and Schaefer atlases.

**Figure S2.**
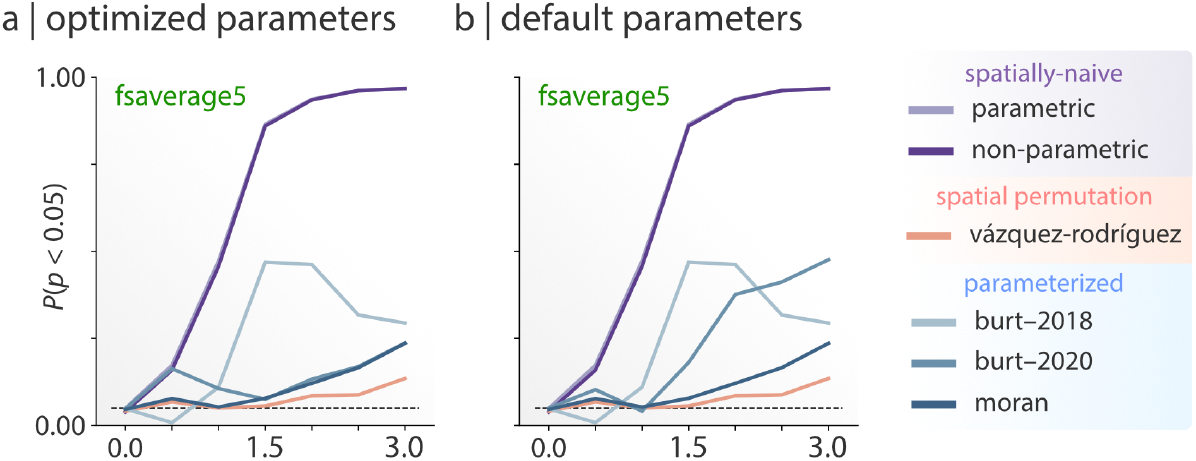
Hyperparameter selection impacts *Burt-2020* model performance. False positive rates of the *Burt-2020* null framework when using optimized (a) versus default (b) settings for the knn hyperparameter in the BrainSMASH software. Here, “optimized” indicates setting the knn parameter equal to the number of vertices in the input data (9,354 and 9,361 for the left and right hemispheres of the fsaverage5 surface; the default setting is 1,000). Optimality was determined by visually evaluating the fit of the generated surrogates as suggested by the authors of the software. Note that all other null framework results are identical between the two figure panels.

**Figure S3.**
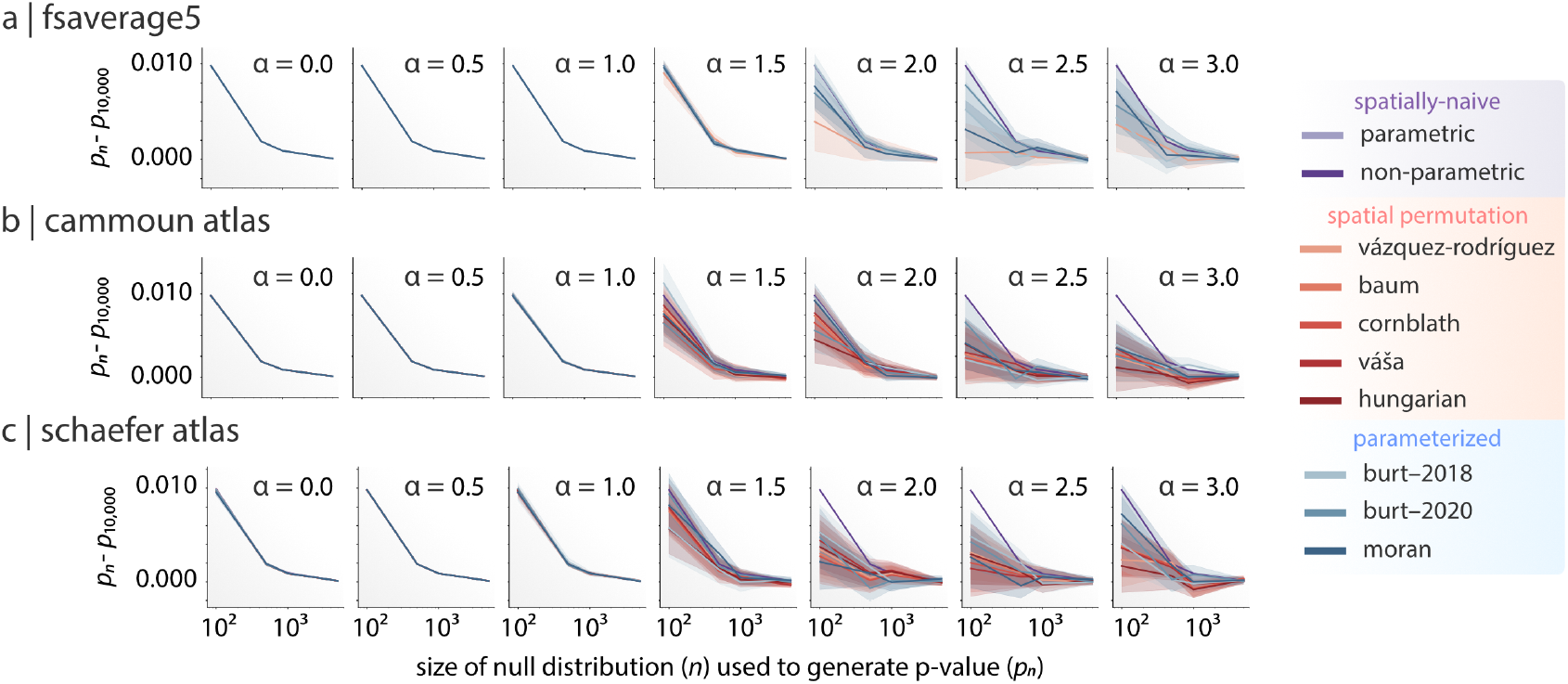
Null models converge rapidly as a function of null distribution size. Difference in *p*-values for each of the null frameworks as a function of the size of the null distribution used to calculate them. Each null framework was used to calculate a “baseline” *p*-value, derived from a distribution of 10,000 null maps (*p*_10,000_). Subsamples of size *n* (where *n* ∈ {100, 500, 1000, 5000}) were drawn from this distribution 1,000 times, and the resulting change in *p*-value was calculated as |*pn − p*_10,000_|. Results are shown for the fsaverage5 surface (a) and the 1,000-parcel Cammoun (b), and 1,000-parcel Schaefer (c) atlases. Colored lines and shaded regions on each plot represent the mean and 95% confidence interval.

**Figure S4.**
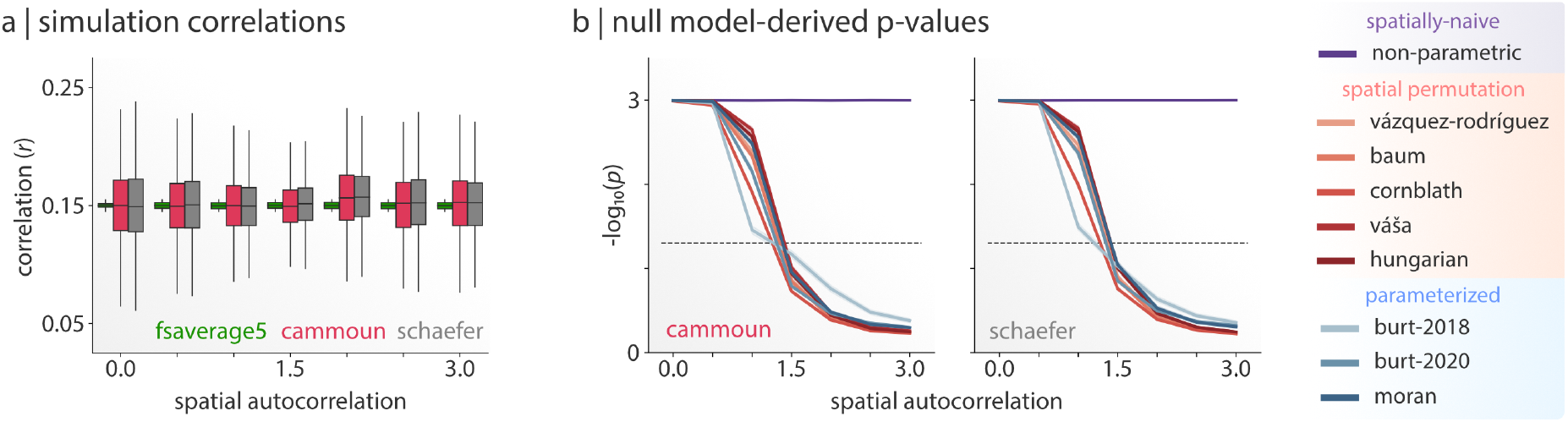
Parcellating simulated data modifies correlation structure. (a) Correlations between pairs of brain maps for all simulations for the vertex (fsaverage5) data—experimentally restricted to the 0.145-0.155 range—and parcellated data across all levels of spatial autocorrelation. Note that parcellating the data increases the range of correlations. (b) Average null model performance for parcellated simulations across all seven levels of spatial autocorrelation, restricting analyses to those simulations where the correlation between brain map pairs was in the range 0.125–0.175. Colored lines and shaded regions on each plot represent the mean and 95% confidence interval. The Cammoun and Schaefer atlas results are shown for the highest resolution (1,000 parcels) only. The dashed black line corresponds to *p* = 0.05 (where values beneath the line indicate *p* > 0.05). Note that the spatially-naive parametric results are not depicted as they are largely indistinguishable from *p* = 0 and approach infinity on the provided scale.

**Figure S5.**
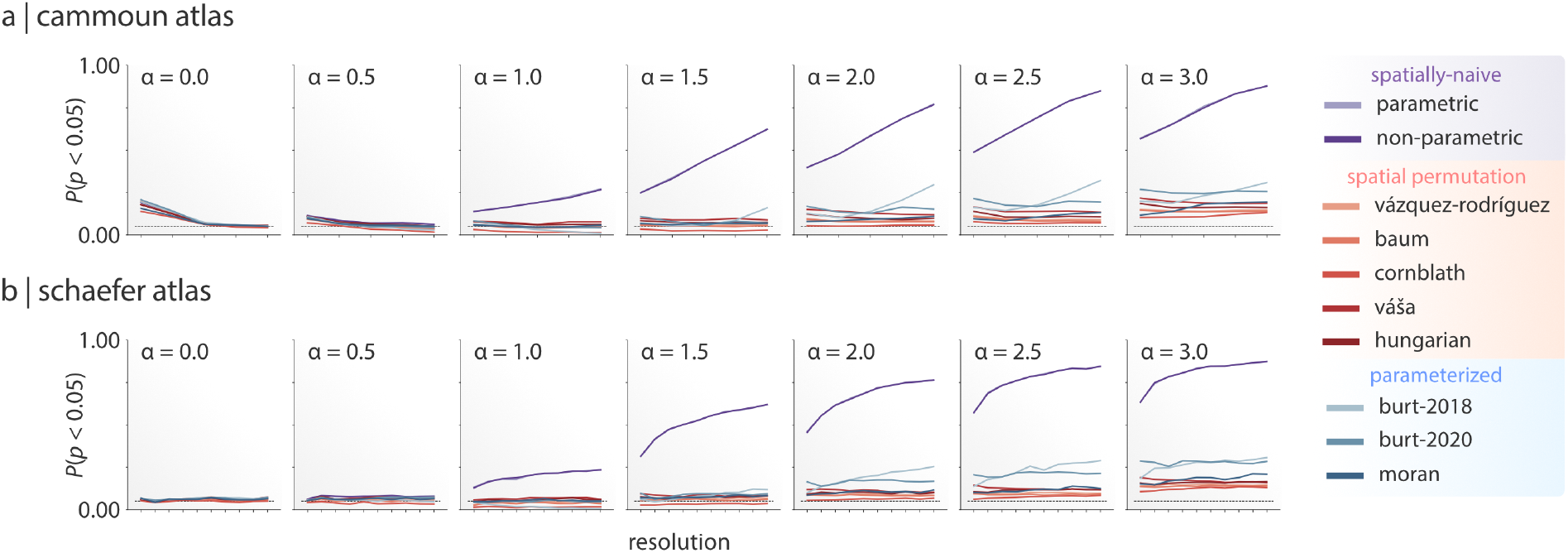
Data resolution does not impact false positive rate of null frameworks. False positive rates for the Cammoun (a) and Schaefer (b) atlases as a function of parcellation resolution for the different null frameworks across all levels of spatial autocorrelation. The dashed black line corresponds to the expected FPR of five percent. Note that the spatially-naive null models (*parametric* and *non-parametric*) are almost completely overlapping on these plots.

**Figure S6.**
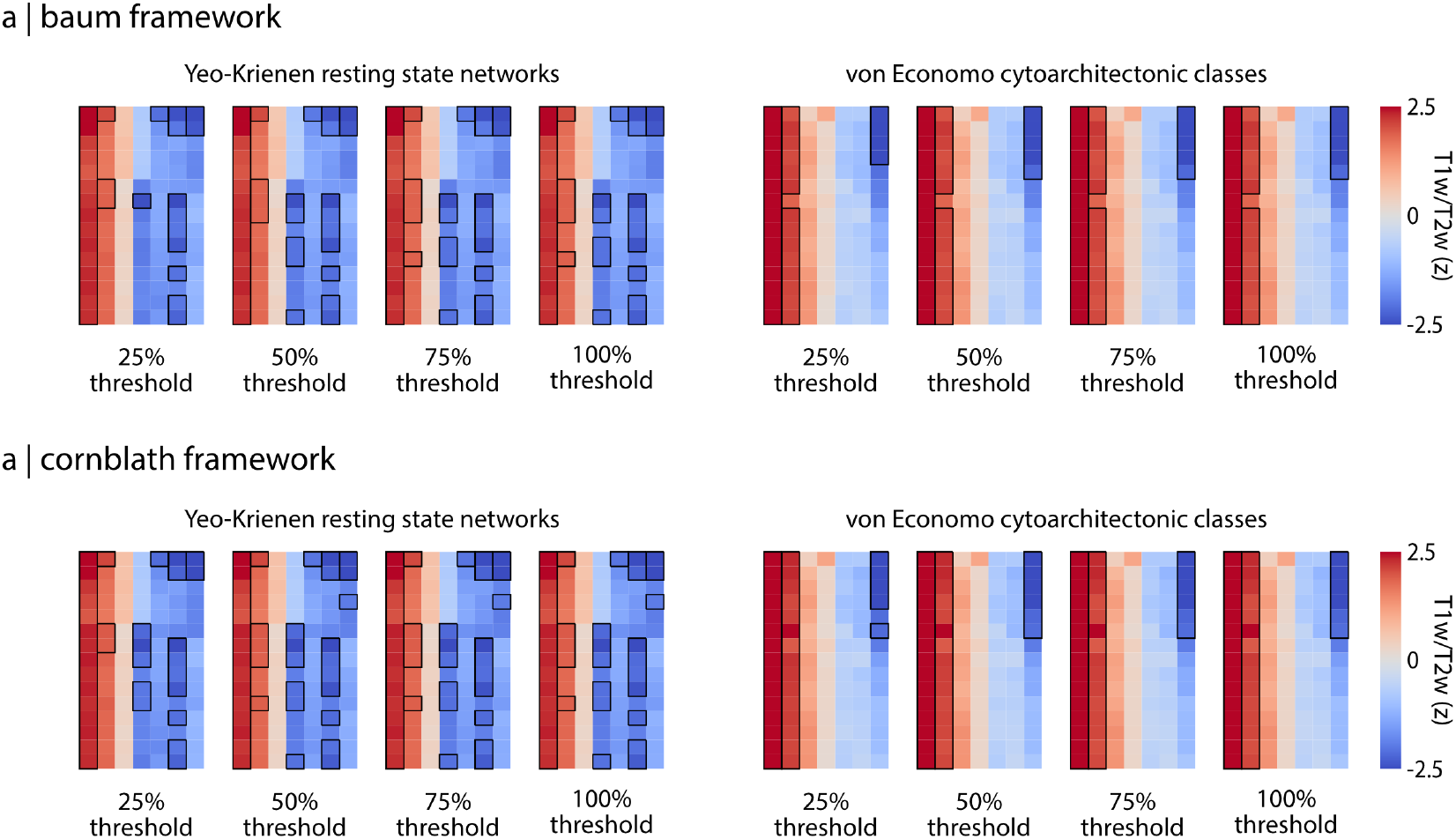
Partition specificity results are not influenced by medial wall rotations. Examination of how constraining the *Baum* (a) and *Cornblath* (b) null methods impacts results of the partition specificity analysis. The analysis procedure was modified such that we removed spatial rotations wherein a given partition had more than a set percent (X%) of all its constituent parcels discarded due to rotation of the medial wall. Each cell in the heatmap represents the T1w/T2w z-score for a given network (x-axis) and atlas / resolution (y-axis). Black outlines around a cell represent network z-scores that are significant at the *p* ≤ 0.05 level. Refer to Fig. 6 for a more detailed overview of the heatmaps. Note that the 100% threshold heatmaps are identical to those in Fig. 6.

**Figure S7.**
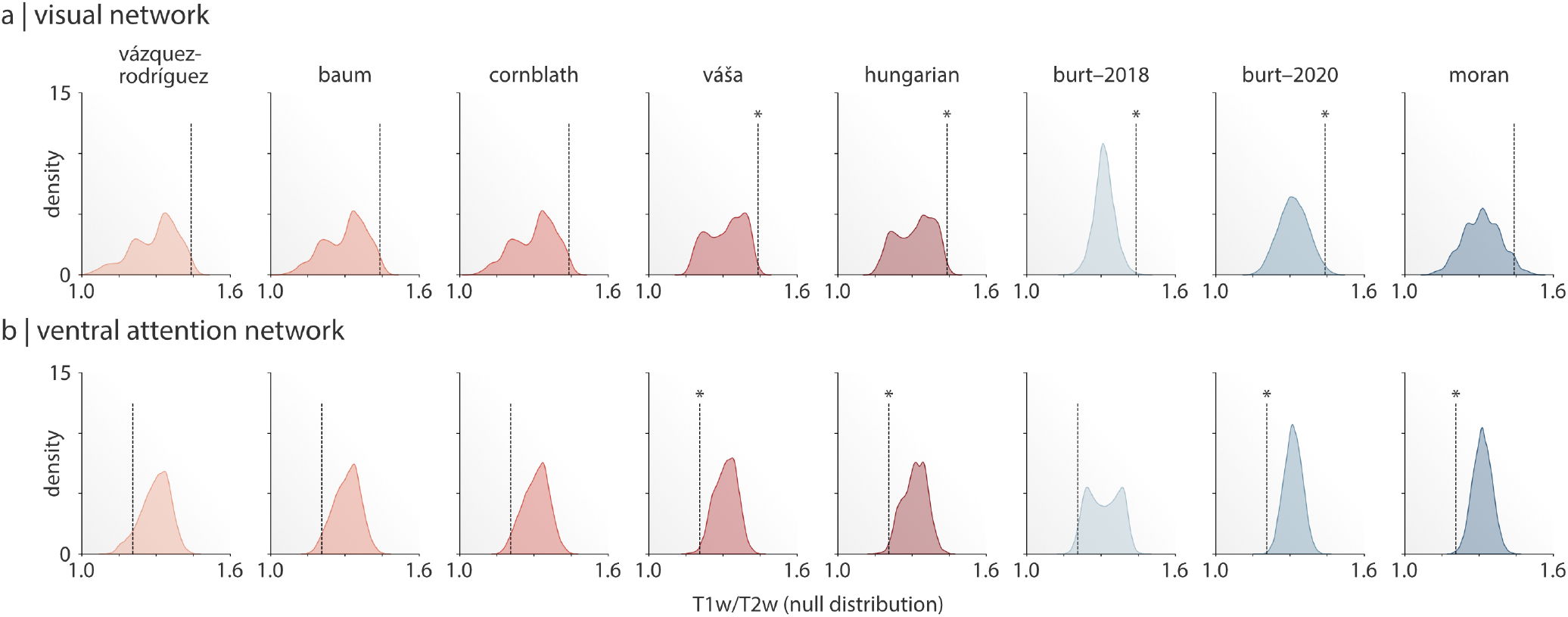
Null distribution shape impacts significance testing. Null distributions of the T1w/T2w ratio for the visual (a) and ventral attention (b) networks from the partition specificity analysis (*Results: Testing partition specificity*), shown here for the 1,000-parcel Cammoun atlas. Dashed lines denote the empirical T1w/T2w value for the network, and an asterisk denotes that the null model yielded a significant test for over-/under-expression of T1w/T2w in the network. Note the similarity in shape of the null distributions for the *Vázquez-Rodríguez*, *Baum*, and *Cornblath* methods and for the *Váša* and *Hungarian* methods.

**Figure S8.**
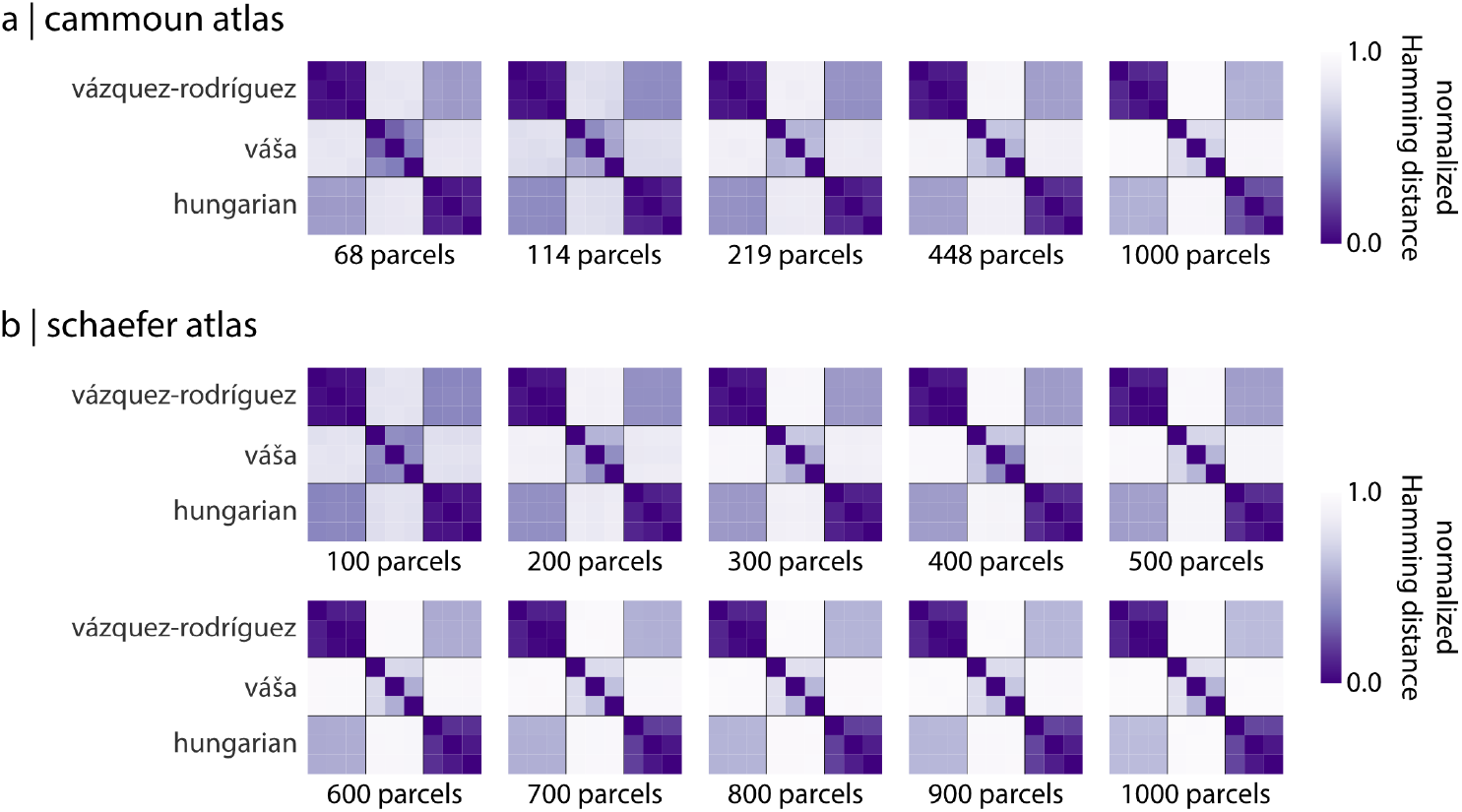
Variability in parcel centroid definition. Reproduction of results shown in Fig. 7a for all resolutions of the (a) Cammoun and (b) Schaefer atlases. Cells within each section of the heatmap represent different parcel centroid definition methods (*surface*, *average*, *geodesic* from top-to-bottom). Refer to *Methods: Null model implementation variability* and Fig. 7 for more information

## References

Akbarian, S., Liu, C., Knowles, J. A., Vaccarino, F. M., Farnham, P. J., Crawford, G. E., Jaffe, A. E., Pinto, D., Dracheva, S., Geschwind, D. H., Mill, J., Nairn, A. C., Abyzov, A., Pochareddy, S., Prabhakar, S., Weissman, S., Sullivan, P. F., State, M. W., Weng, Z., Peters, M. A., White, K. P., Gerstein, M. B., Amiri, A., Armoskus, C., Ashley-Koch, A. E., Bae, T., Beckel-Mitchener, A., Berman, B. P., Coetzee, G. A., Coppola, G., Francoeur, N., Fromer, M., Gao, R., Grennan, K., Herstein, J., Kavanagh, D. H., Ivanov, N. A., Jiang, Y., Kitchen, R. R., Kozlenkov, A., Kundakovic, M., Li, M., Li, Z., Liu, S., Mangravite, L. M., Mattei, E., Markenscoff-Papadimitriou, E., Navarro, F. C. P., North, N., Omberg, L., Panchision, D., Parikshak, N., Poschmann, J., Price, A. J., Purcaro, M., Reddy, T. E., Roussos, P., Schreiner, S., Scuderi, S., Sebra, R., Shibata, M., Shieh, A. W., Skarica, M., Sun, W., Swarup, V., Thomas, A., Tsuji, J., van Bakel, H., Wang, D., Wang, Y., Wang, K., Werling, D. M., Willsey, A. J., Witt, H., Won, H., Wong, C. C. Y., Wray, G. A., Wu, E. Y., Xu, X., Yao, L., Senthil, G., Lehner, T., Sklar, P., and Sestan, N. (2015). The PsychENCODE project. Nature Neuroscience, 18(12):1707–1712.

Alexander-Bloch, A., Raznahan, A., Bullmore, E., and Giedd, J. (2013). The convergence of maturational change and structural covariance in human cortical networks. Journal of Neuroscience, 33(7):2889–2899.

Alexander-Bloch, A. F., Shou, H., Liu, S., Satterthwaite, T. D., Glahn, D. C., Shinohara, R. T., Vandekar, S. N., and Raznahan, (2018). On testing for spatial correspondence between maps of human brain structure and function. NeuroImage, 178:540–551.

Arslan, S., Ktena, S. I., Makropoulos, A., Robinson, E. C., Rueckert, D., and Parisot, S. (2018). Human brain mapping: A systematic comparison of parcellation methods for the human cerebral cortex. Neuroimage, 170:5–30.

Baum, G. L., Cui, Z., Roalf, D. R., Ciric, R., Betzel, R. F., Larsen,, Cieslak, M., Cook, P. A., Xia, C. H., Moore, T. M., Ruparel, K., Oathes, D. J., Alexander-Bloch, A. F., Shinohara, R. T., Raznahan, A., Gur, R. E., Gur, R. C., Bassett, D. S., and Satterthwaite, T. D. (2020). Development of structure–function coupling in human brain networks during youth. Proceedings of the National Academy of Sciences, 117(1):771–778.

Beliveau, V., Ganz, M., Feng, L., Ozenne, B., Højgaard, L., Fisher, P. M., Svarer, C., Greve, D. N., and Knudsen, G. M. (2017). A high-resolution in vivo atlas of the human brain’s serotonin system. Journal of Neuroscience, 37(1):120–128.

Bellec, P., Rosa-Neto, P., Lyttelton, O. C., Benali, H., and Evans, A. C. (2010). Multi-level bootstrap analysis of stable clusters in resting-state fMRI. Neuroimage, 51(3):1126–1139.

Bhagwat, N., Barry, A., Dickie, E. W., Brown, S. T., Devenyi, G. A., Hatano, K., DuPre, E., Dagher, A., Chakravarty, M. M., Greenwood, C. M., Mišić, B., Kennedy, D. N., and Poline, J.-B. (2020). Understanding the impact of preprocessing pipelines on neuroimaging cortical surface analyses. bioRxiv.

Botvinik-Nezer, R., Holzmeister, F., Camerer, C. F., Dreber, A., Huber, J., Johannesson, M., Kirchler, M., Iwanir, R., Mumford, J. A., Adcock, A., Avesani, P., Baczkowski, B., Bajracharya, A., Bakst, L., Ball, S., Barilari, M., Bault, N., Beaton, D., Beitner, J., Benoit, R., Berkers, R., Bhanji, J., Biswal, B., Bobadilla-Suarez, S., Bortolini, T., Bottenhorn, K., Bowring, A., Braem, S., Brooks, H., Brudner, E., Calderon, C., Camilleri, J., Castrellon, J., Cecchetti, L., Cieslik, E., Cole, Z., Collignon, O., Cox, R., Cunningham, W., Czoschke, S., Dadi, K., Davis, C., De Luca, A., Delgado, M., Demetriou, L., Dennison, J., Di, X., Dickie, E., Dobryakova, E., Donnat, C., Dukart, J., Duncan, N. W., Durnez, J., Eed, A., Eickhoff, S., Erhart, A., Fontanesi, L., Fricke, G. M., Galvan, A., Gau, R., Genon, S., Glatard, T., Glerean, E., Goeman, J., Golowin, S., González-García, C., Gorgolewski, K., Grady, C., Green, M., Guassi Moreira, J., Guest, O., Hakimi, S., Hamilton, J. P., Hancock, R., Handjaras, G., Harry, B., Hawco, C., Herholz, P., Herman, G., Heunis, S., Hoffstaedter, F., Hogeveen, J., Holmes, S., Hu, C.-P., Huettel, S., Hughes, M., Iacovella, V., Iordan, A., Isager, P., Isik, A. I., Jahn, A., Johnson, M., Johnstone, T., Joseph, M., Juliano, A., Kable, J., Kassinopoulos, M., Koba, C., Kong, X.-Z., Koscik, T., Kucukboyaci, N. E., Kuhl, B., Kupek, S., Laird, A., Lamm, C., Langner, R., Lauharatanahirun, N., Lee, H., Lee, S., Leemans, A., Leo, A., Lesage, E., Li, F., Li, M., Lim, P. C., Lintz, E., Liphardt, S., Losecaat Vermeer, A., Love, B., Mack, M., Malpica, N., Marins, T., Maumet, C., McDonald, K., McGuire, J., Melero, H., Méndez Leal, A., Meyer, B., Meyer, K., Mihai, P., Mitsis, G., Moll, J., Nielson, D., Nilsonne, G., Notter, M., Olivetti, E., Onicas, A., Papale, P., Patil, K., Peelle, J. E., Pérez, A., Pischedda, D., Poline, J.-B., Prystauka, Y., Ray, S., Reuter-Lorenz, P., Reynolds, R., Ricciardi, E., Rieck, J., Rodriguez-Thompson, A., Romyn, A., Salo, T., Samanez-Larkin, G., Sanz-Morales, E., Schlichting, M., Schultz, D., Shen, Q., Sheridan, M., Shiguang, F., Silvers, J., Skagerlund, K., Smith, A., Smith, D., Sokol-Hessner, P., Steinkamp, S., Tashjian, S., Thirion, B., Thorp, J., Tinghög, G., Tisdall, L., Tompson, S., Toro-Serey, C., Torre, J., Tozzi, L., Truong, V., Turella, L., van’t Veer, A. E., Verguts, T., Vettel, J., Vijayarajah, S., Vo, K., Wall, M., Weeda, W. D., Weis, S., White, D., Wisniewski, D., Xifra-Porxas, A., Yearling, E., Yoon, S., Yuan, R., Yuen, K., Zhang, L., Zhang, X., Zosky, J., Nichols, T. E., Poldrack, R. A., and Schonberg, T. (2020). Variability in the analysis of a single neuroimaging dataset by many teams. Nature.

Breakspear, M., Brammer, M., and Robinson, P. A. (2003). Construction of multivariate surrogate sets from nonlinear data using the wavelet transform. Physica D: Nonlinear Phenomena, 182(1-2):1–22.

Breakspear, M., Brammer, M. J., Bullmore, E. T., Das, P., and Williams, L. M. (2004). Spatiotemporal wavelet resampling for functional neuroimaging data. Human Brain Mapping, 23(1):1–25.

Brett, M., Markiewicz, C. J., Hanke, M., Côté, M.-A., Cipollini, B., McCarthy, P., Jarecka, D., Cheng, C. P., Halchenko, Y. O., Cottaar, M., Ghosh, S., Larson, E., Wassermann, D., Gerhard, S., Lee, G. R., Wang, H.-T., Kastman, E., Kaczmarzyk, J., Guidotti, R., Duek, O., Rokem, A., Madison, C., Morency, F. C., Moloney, B., Goncalves, M., Markello, R., Riddell, C., Burns, C., Millman, J., Gramfort, A., Leppäkangas, J., Sólon, A., van den Bosch, J. J., Vincent, R. D., Braun, H., Subramaniam, K., Gorgolewski, K. J., Raamana, P. R., Nichols, B. N., Baker, E. M., Hayashi, S., Pinsard, B., Haselgrove, C., Hymers, M., Esteban, O., Koudoro, S., Oosterhof, N. N., Amirbekian, B., Nimmo-Smith, I., Nguyen, L., Reddigari, S., St-Jean, S., Panfilov, E., Garyfallidis, E., Varoquaux, G., Legarreta, J. H., Hahn, K. S., Hinds, O. P., Fauber, B., Poline, J.-B., Stutters, J., Jordan, K., Cieslak, M., Moreno, M. E., Haenel, V., Schwartz, Y., Baratz, Z., Darwin, B. C., Thirion, B., Papadopoulos Orfanos, D., Pérez-García, F., Solovey, I., Gonzalez, I., Palasubramaniam, J., Lecher, J., Leinweber, K., Raktivan, K., Fischer, P., Gervais, P., Gadde, S., Ballinger, T., Roos, T., and Reddam, V. R. (2020). nipy/nibabel. Zenodo, doi:10.5281/zenodo.591597.

Burt, J. B., Demirtaş, M., Eckner, W. J., Navejar, N. M., Ji, J. L., Martin, W. J., Bernacchia, A., Anticevic, A., and Murray, J. D. (2018). Hierarchy of transcriptomic specialization across human cortex captured by structural neuroimaging topography. Nature Neuroscience, 21(9):1251–1259.

Burt, J. B., Helmer, M., Shinn, M., Anticevic, A., and Murray, J. D. (2020). Generative modeling of brain maps with spatial autocorrelation. NeuroImage.

Cammoun, L., Gigandet, X., Meskaldji, D., Thiran, J. P., Sporns, O., Do, K. Q., Maeder, P., Meuli, R., and Hagmann, P. (2012). Mapping the human connectome at multiple scales with diffusion spectrum MRI. Journal of Neuroscience Methods, 203(2):386–397.

Carp, J. (2012). On the plurality of (methodological) worlds: estimating the analytic flexibility of fMRI experiments. Frontiers in Neuroscience, 6:149.

Casey, B. J., Cannonier, T., Conley, M. I., Cohen, A. O., Barch, D. M., Heitzeg, M. M., Soules, M. E., Teslovich, T., Dellarco, D. V., Garavan, H., Orr, C. A., Wager, T. D., Banich, M. T., Speer, N. K., Sutherland, M. T., Riedel, M. C., Dick, A. S., Bjork, J. M., Thomas, K. M., Chaarani, B., Mejia, M. H., Jr., D. J. H., Cornejo, M. D., Sicat, C. S., Harms, M. P., Dosenbach, N. U. F., Rosenberg, M., Earl, E., Bartsch, H., Watts, R., Polimeni, J. R., Kuperman, J. M., Fair, D. A., Dale, A. M., and the ABCD Imaging Acquisition Workgroup (2018). The Adolescent Brain Cognitive Development (ABCD) study: Imaging acquisition across 21 sites. Developmental Cognitive Neuroscience, 32:43–54.

Ciric, R., Wolf, D. H., Power, J. D., Roalf, D. R., Baum, G. L., Ruparel, K., Shinohara, R. T., Elliott, M. A., Eickhoff, S. B., Davatzikos, C., Gur, R. C., Gur, R. E., Bassett, D. S., and Satterthwaite, T. D. (2017). Benchmarking of participant-level confound regression strategies for the control of motion artifact in studies of functional connectivity. Neuroimage, 154:174–187.

Cliff, A. D. and Ord, K. (1970). Spatial autocorrelation: a review of existing and new measures with applications. Economic Geography, 46(sup1):269–292.

Cornblath, E. J., Ashourvan, A., Kim, J. Z., Betzel, R. F., Ciric, R., Adebimpe, A., Baum, G. L., He, X., Ruparel, K., Moore, T. M., Gur, R. C., Gur, R. E., Shinohara, R. T., Roalf, D. R., Satterthwaite, T. D., and Bassett, D. S. (2020). Temporal sequences of brain activity at rest are constrained by white matter structure and modulated by cognitive demands. Communications Biology, 3(1):1–12.

Cressie, N. (2015). Statistics for spatial data. John Wiley & Sons.

Dadi, K., Varoquaux, G., Machlouzarides-Shalit, A., Gorgolewski, K. J., Wassermann, D., Thirion, B., and Mensch, A. (2020). Fine-grain atlases of functional modes for fMRI analysis. NeuroImage.

Damoiseaux, J. S., Rombouts, S., Barkhof, F., Scheltens, P., Stam, C. J., Smith, S. M., and Beckmann, C. F. (2006). Consistent resting-state networks across healthy subjects. Proceedings of the National Academy of Sciences, 103(37):13848–13853.

Deblauwe, V., Kennel, P., and Couteron, P. (2012). Testing pairwise association between spatially autocorrelated variables: A new approach using surrogate lattice data. PloS One, 7(11):e48766.

Demirtaş, M., Burt, J. B., Helmer, M., Ji, J. L., Adkinson, B. D., Glasser, M. F., Van Essen, D. C., Sotiropoulos, S. N., Anticevic, A., and Murray, J. D. (2019). Hierarchical heterogeneity across human cortex shapes large-scale neural dynamics. Neuron, 101(6):1181–1194.

Desikan, R. S., Ségonne, F., Fischl, B., Quinn, B. T., Dickerson, B. C., Blacker, D., Buckner, R. L., Dale, A. M., Maguire, R. P., Hyman, B. T., Albert, M. S., and Killiany, R. J. (2006). An automated labeling system for subdividing the human cerebral cortex on MRI scans into gyral based regions of interest. NeuroImage, 31(3):968–980.

Dray, S. (2011). A new perspective about Moran’s coefficient: Spatial autocorrelation as a linear regression problem. Geographical Analysis, 43(2):127–141.

Dutilleul, P., Clifford, P., Richardson, S., and Hemon, D. (1993). Modifying the t test for assessing the correlation between two spatial processes. Biometrics, pages 305–314.

Eickhoff, S. B., Yeo, B. T., and Genon, S. (2018). Imaging-based parcellations of the human brain. Nature Reviews Neuroscience, 19(11):672–686.

Finn, E. S., Shen, X., Scheinost, D., Rosenberg, M. D., Huang, J., Chun, M. M., Papademetris, X., and Constable, R. T. (2015). Functional connectome fingerprinting: identifying individuals using patterns of brain connectivity. Nature Neuroscience, 18(11):1664–1671.

Fischl, B., Sereno, M. I., Tootell, R. B., and Dale, A. M. (1999). High-resolution intersubject averaging and a coordinate system for the cortical surface. Human Brain Mapping, 8(4):272–284.

Fortin, M.-J. and Jacquez, G. M. (2000). Randomization tests and spatially auto-correlated data. Bulletin of the Ecological Society of America, 81(3):201–205.

Fulcher, B. D., Arnatkeviciute, A., and Fornito, A. (2020). Overcoming bias in gene-set enrichment analyses of brain-wide transcriptomic data. bioRxiv.

Gao, R., van den Brink, R. L., Pfeffer, T., and Voytek, B. (2020). Neuronal timescales are functionally dynamic and shaped by cortical microarchitecture. bioRxiv.

Glasser, M. F. and Van Essen, D. C. (2011). Mapping human cortical areas in vivo based on myelin content as revealed by T1-and T2-weighted MRI. Journal of Neuroscience, 31(32):11597–11616.

Gordon, E. M., Laumann, T. O., Adeyemo, B., Huckins, J. F., Kelley, W. M., and Petersen, S. E. (2016). Generation and evaluation of a cortical area parcellation from resting-state correlations. Cerebral cortex, 26(1):288–303.

Gorgolewski, K. J., Varoquaux, G., Rivera, G., Schwarz, Y., Ghosh, S. S., Maumet, C., Sochat, V. V., Nichols, T. E., Poldrack, R. A., Poline, J.-B., Yarkoni, T., and Margulies, D. S. (2015). Neurovault.org: a web-based repository for collecting and sharing unthresholded statistical maps of the human brain. Frontiers in Neuroinformatics, 9:8.

Hamming, R. W. (1950). Error detecting and error correcting codes. The Bell System Technical Journal, 29(2):147–160.

Hansen, J. Y., Markello, R. D., Vogel, J. W., Seidlitz, J., Bzdok, D., and Mišić, B. (2020). Molecular signatures of cognition and affect. bioRxiv.

Hawrylycz, M. J., Lein, E. S., Guillozet-Bongaarts, A. L., Shen, E. H., Ng, L., Miller, J. A., van de Lagemaat, L. N., Smith, K. A., Ebbert, A., Riley, Z. L., Abajian, C., Beckmann, C. F., Bernard, A., Bertagnolli, D., Boe, A. F., Cartagena, P. M., Chakravarty, M. M., Chapin, M., Chong, J., Dalley, R. A., Daly, B. D., Dang, C., Datta, S., Dee, N., Dolbeare, T. A., Faber, V., Feng, D., Fowler, D. R., Goldy, J., Gregor, B. W., Haradon, Z., Haynor, D. R., Hohmann, J. G., Horvath, S., Howard, R. E., Jeromin, A., Jochim, J. M., Kinnunen, M., Lau, C., Lazarz, E. T., Lee, C., Lemon, T. A., Li, L., Li, Y., Morris, J. A., Overly, C. C., Parker, P. D., Parry, S. E., Reding, M., Royall, J. J., Schulkin, J., Sequeira, P. A., Slaughterbeck, C. R., Smith, S. C., Sodt, A. J., Sunkin, S. M., Swanson, B. E., Vawter, M. P., Williams, D., Wohnoutka, P., Zielke, H. R., Geschwind, D. H., Hof, P. R., Smith, S. M., Koch, C., Grant, S. G. N., and Jones, A. R. (2012). An anatomically comprehensive atlas of the adult human brain transcriptome. Nature, 489(7416):391.

Hunter, J. D. (2007). Matplotlib: A 2D graphics environment. Computing in Science & Engineering, 9(3):90–95.

Insel, T. R., Landis, S. C., and Collins, F. S. (2013). The NIH BRAIN initiative. Science, 340(6133):687–688.

Kharabian Masouleh, S., Eickhoff, S., Zeighami, Y., Lewis, L., Dahnke, R., Gaser, R., Gaser, C., Chouinard-Decorte, F., Lepage, C., Scholtens, L., Hoffstaedter, F., Glahn, D., Blangero, J., Evans, A., Genon, S., and Valk, S. L. (2020). Influence of processing pipeline on cortical thickness measurement. Cerebral Cortex.

Kluyver, T., Ragan-Kelley, B., Pérez, F., Granger, B., Bussonnier, M., Frederic, J., Kelley, K., Hamrick, J., Grout, J., Corlay, S., Ivanov, P., Avila, D., Abdalla, S., Willing, C., and the Jupyter development team (2016). Jupyter Notebooks–A publishing format for reproducible computational workflows. In Loizides, F. and Scmidt, B., editors, Positioning and Power in Academic Publishing: Players, Agents and Agendas, pages 87–90. IOS Press.

Kong, R., Li, J., Orban, C., Sabuncu, M. R., Liu, H., Schaefer, A., Sun, N., Zuo, X.-N., Holmes, A. J., Eickhoff, S. B., and Yeo, B. T. T. (2019). Spatial topography of individual-specific cortical networks predicts human cognition, personality, and emotion. Cerebral Cortex, 29(6):2533–2551.

Kuhn, H. W. (1955). The hungarian method for the assignment problem. Naval research logistics quarterly, 2(1-2):83–97.

Legendre, P. (1993). Spatial autocorrelation: trouble or new paradigm? Ecology, 74(6):1659–1673.

Maier-Hein, K. H., Neher, P. F., Houde, J.-C., Côté, M.-A., Gary-fallidis, E., Zhong, J., Chamberland, M., Yeh, F.-C., Lin, Y.-C., Ji, Q., Reddick, W. E., Glass, J. O., Chen, D. Q., Feng, Y., Gao, C., Wu, Y., Ma, J., He, R., Li, Q., Westin, C.-F., Deslauriers-Gauthier, S., González, J. O. O., Paquette, M., St-Jean, S., Girard, G., Rheault, F., Sidhu, J., Tax, C. M. W., Guo, F., Mesri, H. Y., Dávid, S., Froeling, M., Heemskerk, A. M., Leemans, A., Boré, A., Pinsard, B., Bedetti, C., Desrosiers, M., Brambati, S., Doyon, J., Sarica, A., Vasta, R., Cerasa, A., Quattrone, A., Yeatman, J., Khan, A. R., Hodges, W., Alexander, S., Romascano, D., Barakovic, M., Auría, A., Esteban, O., Lemkaddem, A., Thiran, J.-P., Cetingul, H. E., Odry, B. L., Mailhe, B., Nadar, M. S., Pizzagalli, F., Prasad, G., Villalon-Reina, J. E., Galvis, J., Thompson, P. M., Requejo, F. D. S., Laguna, P. L., Lacerda, L. M., Barrett, R., Dell’Acqua, F., Catani, M., Petit, L., Caruyer, E., Daducci, A., Dyrby, T. B., Holland-Letz, T., Hilgetag, C. C., Stieltjes, B., and Descoteaux, M. (2017). The challenge of mapping the human connectome based on diffusion tractography. Nature Communications, 8(1):1–13.

Marcus, D., Harwell, J., Olsen, T., Hodge, M., Glasser, M., Prior, F., Jenkinson, M., Laumann, T., Curtiss, S., and Van Essen, D. (2011). Informatics and data mining tools and strategies for the human connectome project. Frontiers in Neuroinformatics, 5:4.

Margulies, D. S., Ghosh, S. S., Goulas, A., Falkiewicz, M., Huntenburg, J. M., Langs, G., Bezgin, G., Eickhoff, S. B., Castellanos, F. X., Petrides, M., Jeffries, E., and Smallwood, J. (2016). Situating the default-mode network along a principal gradient of macroscale cortical organization. Proceedings of the National Academy of Sciences, 113(44):12574–12579.

McKinney, W. (2010). Data structures for statistical computing in Python. In Proceedings of the 9th Python in Science Conference, volume 445, pages 51–56. Austin, TX.

Messé, A. (2020). Parcellation influence on the connectivity-based structure–function relationship in the human brain. Human Brain Mapping, 41(5):1167–1180.

Murray, J. D., Bernacchia, A., Freedman, D. J., Romo, R., Wallis, J. D., Cai, X., Padoa-Schioppa, C., Pasternak, T., Seo, H., Lee, D., and Wang, X.-J. (2014). A hierarchy of intrinsic timescales across primate cortex. Nature Neuroscience, 17(12):1661–1663.

Norgaard, M., Beliveau, V., Ganz, M., Svarer, C., Pinborg, L. H., Keller, S. H., Jensen, P. S., Greve, D. N., and Knudsen, G. M. (2020). A high-resolution in vivo atlas of the human brain’s benzodiazepine binding site of GABA*A* receptors. bioRxiv.

Oldham, S., Arnatkeviciute, A., Smith, R. E., Tiego, J., Bellgrove, M. A., and Fornito, A. (2020). The efficacy of different preprocessing steps in reducing motion-related confounds in diffusion MRI connectomics. bioRxiv.

Oliphant, T. E. (2006). A guide to NumPy, volume 1. Trelgol Publishing USA.

Paquola, C., Seidlitz, J., Benkarim, O., Royer, J., Klimes, P., Bethlehem, R. A., Larivière, S., de Wael, R. V., Rodríguez-Cruces, R., Hall, J. A., Frauscher, B., Smallwood, J., and Bernhardt, B. C. (2020). A multi-scale cortical wiring space links cellular architecture and functional dynamics in the human brain. PLoS Biology, 18(11):e3000979.

Parkes, L., Fulcher, B., Yücel, M., and Fornito, A. (2018). An evaluation of the efficacy, reliability, and sensitivity of motion correction strategies for resting-state functional mri. NeuroImage, 171:415–436.

Pedregosa, F., Varoquaux, G., Gramfort, A., Michel, V., Thirion, B., Grisel, O., Blondel, M., Prettenhofer, P., Weiss, R., Dubourg, V., Vanderplas, J., Passos, A., Cournapeau, D., Brucher, M., Perrot, M., and Duchesnay, E. (2011). Scikit-learn: Machine learning in Python. Journal of Machine Learning Research, 12(Oct):2825–2830.

Pérez, F. and Granger, B. E. (2007). IPython: A system for interactive scientific computing. Computing in Science & Engineering, 9(3):21–29.

Poldrack, R. A. (2011). Inferring mental states from neuroimaging data: From reverse inference to large-scale decoding. Neuron, 72(5):692–697.

Poldrack, R. A., Barch, D. M., Mitchell, J., Wager, T., Wagner, A. D., Devlin, J. T., Cumba, C., Koyejo, O., and Milham, M. (2013). Toward open sharing of task-based fMRI data: the OpenfMRI project. Frontiers in Neuroinformatics, 7:12.

Poldrack, R. A., Kittur, A., Kalar, D., Miller, E., Seppa, C., Gil, Y., Parker, D. S., Sabb, F. W., and Bilder, R. M. (2011). The cognitive atlas: toward a knowledge foundation for cognitive neuroscience. Frontiers in Neuroinformatics, 5:17.

Poldrack, R. A. and Yarkoni, T. (2016). From brain maps to cognitive ontologies: Informatics and the search for mental structure. Annual Review of Psychology, 67:587–612.

Proix, T., Spiegler, A., Schirner, M., Rothmeier, S., Ritter, P., and Jirsa, V. K. (2016). How do parcellation size and shortrange connectivity affect dynamics in large-scale brain network models? NeuroImage, 142:135–149.

Royer, J., Paquola, C., Larivière, S., de Wael, R. V., Tavakol, S., Lowe, A. J., Benkarim, O., Evans, A. C., Bzdok, D., Smallwood, J., Frauscher, B., and Bernhardt, B. C. (2020). Myeloarchitecture gradients in the human insula: Histological underpinnings and association to intrinsic functional connectivity. NeuroImage, page 116859.

Schaefer, A., Kong, R., Gordon, E. M., Laumann, T. O., Zuo, X.-N., Holmes, A. J., Eickhoff, S. B., and Yeo, B. T. (2018). Local-global parcellation of the human cerebral cortex from intrinsic functional connectivity MRI. Cerebral Cortex, 28(9):3095–3114.

Schilling, K. G., Nath, V., Hansen, C., Parvathaneni, P., Blaber, J., Gao, Y., Neher, P., Aydogan, D. B., Shi, Y., Ocampo-Pineda, M., Schiavi, S., Daducci, A., Girard, G., Barakovic, M., Rafael-Patino, J., Romascano, D., Rensonnet, G., Pizzolato, M., Bates, A., Fischi, E., Thiran, J.-P., Canales-Rodríguez, E. J., Huang, C., Zhu, H., Zhong, L., Cabeen, R., Toga, A. W., Rheault, F., Theaud, G., Houde, J.-C., Sidhu, J., Chamberland, M., Westin, C.-F., Dyrby, T. B., Verma, R., Rathi, Y., Irfanoglu, M. O., Thomas, C., Pierpaoli, C., Descoteaux, M., Anderson, A. W., and Landman, B. A. (2019). Limits to anatomical accuracy of diffusion tractography using modern approaches. NeuroImage, 185:1–11.

Scholtens, L. H., de Reus, M. A., de Lange, S. C., Schmidt, R., and van den Heuvel, M. P. (2018). An MRI Von Economo–Koskinas atlas. NeuroImage, 170:249–256.

Shafiei, G., Markello, R. D., De Wael, R. V., Bernhardt, B. C., Fulcher, B. D., and Misic, B. (2020). Topographic gradients of intrinsic dynamics across neocortex. Elife, 9:e62116.

Sudlow, C., Gallacher, J., Allen, N., Beral, V., Burton, P., Danesh, J., Downey, P., Elliott, P., Green, J., Landray, M., Liu, B., Matthews, P., Ong, G., Pell, J., Silman, A., Young, A., Sprosen, T., Peakman, T., and Collins, R. (2015). UK Biobank: An open access resource for identifying the causes of a wide range of complex diseases of middle and old age. PLoS Medicine, 12(3).

Van Der Walt, S., Colbert, S. C., and Varoquaux, G. (2011). The NumPy array: a structure for efficient numerical computation. Computing in Science & Engineering, 13(2):22.

Van Essen, D. C., Smith, S. M., Barch, D. M., Behrens, T. E., Yacoub, E., Ugurbil, K., and the Wu-Minn HCP Consortium (2013). The WU-Minn human connectome project: an overview. NeuroImage, 80:62–79.

Váša, F., Seidlitz, J., Romero-Garcia, R., Whitaker, K. J., Rosenthal, G., Vértes, P. E., Shinn, M., Alexander-Bloch, A., Fonagy, P., Dolan, R. J., Jones, P. B., Goodyer, I. M., the NSPN consortium, Sporns, O., and Bullmore, E. T. (2018). Adolescent tuning of association cortex in human structural brain networks. Cerebral Cortex, 28(1):281–294.

Vázquez-Rodríguez, B., Suárez, L. E., Markello, R. D., Shafiei, G., Paquola, C., Hagmann, P., Van Den Heuvel, M. P., Bernhardt, B. C., Spreng, R. N., and Mišić, B. (2019). Gradients of structure–function tethering across neocortex. Proceedings of the National Academy of Sciences, 116(42):21219–21227.

Virtanen, P., Gommers, R., Oliphant, T. E., Haberland, M., Reddy, T., Cournapeau, D., Burovski, E., Peterson, P., Weckesser, W., Bright, J., van der Walt, S. J., Brett, M., Wilson, J., Millman, K. J., Mayorov, N., Nelson, A. R. J., Jones, E., Kern, R., Larson, E., Carey, C. J., İlhan Polat, Feng, Y., Moore, E. W., VanderPlas, J., Laxalde, D., Perktold, J., Cimrman, R., Henriksen, I., Quintero, E. A., Harris, C. R., Archibald, A. M., Ribeiro, A. H., Pedregosa, F., van Mulbregt, P., and the SciPy 1.0 Contributors (2020). Scipy 1.0: fundamental algorithms for scientific computing in python. Nature Methods, pages 1–12.

von Economo, C. F. and Koskinas, G. N. (1925). Die cytoarchitektonik der hirnrinde des erwachsenen menschen. J. Springer.

Vos de Wael, R., Benkarim, O., Paquola, C., Lariviere, S., Royer, J., Tavakol, S., Xu, T., Hong, S.-J., Langs, G., Valk, S., Mišić, B., Milham, M., Margulies, D., Smallwood, J., and Bernhardt, B. C. (2020). BrainSpace: a toolbox for the analysis of macroscale gradients in neuroimaging and connectomics datasets. Communications Biology, 3(1):1–10.

Wagner, H. H. and Dray, S. (2015). Generating spatially constrained null models for irregularly spaced data using Moran spectral randomization methods. Methods in Ecology and Evolution, 6(10):1169–1178.

Wang, P., Kong, R., Kong, X., Liégeois, R., Orban, C., Deco, G., Van Den Heuvel, M. P., and Yeo, B. T. (2019). Inversion of a large-scale circuit model reveals a cortical hierarchy in the dynamic resting human brain. Science Advances, 5(1):eaat7854.

Waskom, M., Botvinnik, O., Ostblom, J., Gelbart, M., Lukauskas, S., Hobson, P., Gemperline, D. C., Augspurger, T., Halchenko, Y., Cole, J. B., Warmenhoven, J., de Ruiter, J., Pye, C., Hoyer, S., Vanderplas, J., Villalba, S., Kunter, G., Quintero, E., Bachant, P., Martin, M., Meyer, K., Swain, C., Miles, A., Brunner, T., O’Kane, D., Yarkoni, T., Williams, M. L., Evans, C., and Fitzgerald, C. (2020a). mwaskom/seaborn. Zenodo, doi:10.5281/zenodo.592845.

Waskom, M., Larson, E., Brodbeck, C., Gramfort, A., Burns, S., Luessi, M., Weidemann, C. T., Bitzer, S., Markiewicz, C., LaPlante, R., Halchenko, Y., Engemann, D. A., van Vliet, M., Ghosh, S., Klein, N., Piantoni, G., Brett, M., Gwilliams, L., King, J.-R., and Liu, D. (2020b). nipy/pysurfer. Zenodo, doi:10.5281/zenodo.592515.

Westfall, P. H. and Young, S. S. (1993). Resampling-based multiple testing: Examples and methods for p-value adjustment, volume 279. John Wiley & Sons.

Whitaker, K. J., Vértes, P. E., Romero-Garcia, R., Váša, F., Moutoussis, M., Prabhu, G., Weiskopf, N., Callaghan, M. F., Wagstyl, K., Rittman, T., Tait, R., Ooi, C., Suckling, J., Inkster, B., Fonagy, P., Dolan, R. J., Jones, P. B., Goodyer, I. M., the NSPN Consortium, and Bullmore, E. T. (2016). Adolescence is associated with genomically patterned consolidation of the hubs of the human brain connectome. Proceedings of the National Academy of Sciences, 113(32):9105–9110.

Witt, S. T., van Ettinger-Veenstra, H., Salo, T., Riedel, M. C., and Laird, A. R. (2020). What executive function network is that? An image-based meta-analysis of network labels. bioRxiv.

Yarkoni, T., Poldrack, R. A., Nichols, T. E., Van Essen, D. C., and Wager, T. D. (2011). Large-scale automated synthesis of human functional neuroimaging data. Nature Methods, 8(8):665.

Yeo, B. T., Krienen, F. M., Sepulcre, J., Sabuncu, M. R., Lashkari, D., Hollinshead, M., Roffman, J. L., Smoller, J. W., Zöllei, L., Polimeni, J. R., Fischl, B., Liu, H., and Buckner, R. L. (2011). The organization of the human cerebral cortex estimated by intrinsic functional connectivity. Journal of Neurophysiology, 106(3):1125.

Zilles, K. and Amunts, K. (2009). Receptor mapping: architecture of the human cerebral cortex. Current Opinion in Neurology, 22(4):331–339.

Zilles, K., Palomero-Gallagher, N., and Schleicher, A. (2004). Transmitter receptors and functional anatomy of the cerebral cortex. Journal of Anatomy, 205(6):417–432.

